# Joint inference and alignment of genome structures enables characterization of compartment-independent 3D relocalization across cell types

**DOI:** 10.1101/545194

**Authors:** Lila Rieber, Shaun Mahony

## Abstract

Cell-type-specific chromosome conformation is correlated with differential gene regulation. We developed MultiMDS to jointly infer and align 3D chromosomal structures from Hi-C datasets, thereby enabling a new way to comprehensively quantify relocalization of genomic loci between cell types. We demonstrate this approach by comparing Hi-C data across a variety of cell types. We consistently find relocalization of loci with minimal difference in A/B compartment score. Some such compartment-independent relocalizations involve loci that display enhancer-associated histone marks in one cell type and polycomb-associated histone marks in the other. MultiMDS thus detects types of relocalizations that are missed by other approaches.

**Availability:** https://github.com/seqcode/multimds

## BACKGROUND

Chromosome conformation data have become available for diverse cell types, perturbations, and developmental stages. Comparing these datasets can highlight the relationship between three-dimensional structure and biological function. To date, such comparisons have primarily examined known chromosomal structures, such as A/B compartments, topologically associated domains (TADs), and loops. Though TADs are largely conserved across cell types and species (1), extensive differences in compartmentalization (2,3) and looping are detectable between datasets (4–6) and are correlated with differential gene expression. However, focusing on known types of differences means that other potentially functional differences remain unexplored. For example, relocalization within a compartment or TAD could be correlated with differences in gene regulation, but could be missed by current approaches for comparing Hi-C datasets.

Current methods for comparing chromosomal conformation datasets (primarily from Hi-C experiments) can be classified as global, interaction-specific, or locus-specific. Some global methods calculate an overall similarity score for two Hi-C datasets, enabling clustering of experiments (7–10). Global concordance in TAD boundaries can also be calculated (11). However, these methods cannot discover specific differences between datasets. Interaction-specific methods identify genomic locus pairs that significantly differ in their interaction frequency, which may indicate gain or loss of a chromatin loop (12–15), but cannot determine how the interacting pair of loci have moved with respect to the rest of the genome. Locus-specific methods identify differences in organization, such as a difference in compartment score or gain or loss of a TAD boundary, that occur at a single locus, which is a single bin at the resolution of the Hi-C data. However, these methods are currently limited to measuring differences in known forms of chromatin organization, preventing the discovery of novel structures. There is currently no method for quantifying general locus-specific relocalizations between Hi-C datasets. Ideally, such a method would quantify the degree to which a given locus has changed position with respect to the rest of the genome, regardless of whether the relocalization was driven by differences in compartmentalization, TAD structure, looping, or a combination of several effects.

In a Hi-C dataset, each locus is represented as a vector of interaction frequencies with every other locus of the genome or chromosome. Because typical metrics for vector comparison, such as Pearson correlation, are biased by Hi-C distance decay, comparison of datasets is challenging (10). To mitigate issues associated with the high dimensionality of Hi-C data, we first aim to embed the datasets in a lower dimensional space. In this work, we choose to embed Hi-C data in three dimensions, representing the population average chromosome structure. While the physical interpretation of 3D chromosome structures is limited by population heterogeneity, we propose that comparing two 3D structures provides a convenient and intuitive assessment of the overall differences in chromatin organization across cell types or conditions. If the structures are comparable and correctly aligned, the degree to which a given locus has shifted position can be calculated as the Euclidean distance between the 3D coordinates of that locus in each of the structures. We have developed MultiMDS to simultaneously infer and align 3D structures from two Hi-C datasets. By applying our method to a number of mammalian and yeast Hi-C datasets, we identified examples of chromatin relocalization correlated with biological function, some of which confirm previous findings and some of which are potentially novel.

## RESULTS

### MultiMDS: a principled approach for comparing genome structures

We developed MultiMDS to quantify locus-specific relocalization between Hi-C datasets. MultiMDS takes as input two Hi-C contact matrices and outputs two aligned 3D structures, which represent the ensemble average structures for the respective inputs (Fig. 1A). Relocalization is calculated as the locus-specific Euclidean distance between aligned structures (Fig. 1B). Because Hi-C contact frequencies are believed to be a function of the average physical distances in the chromosomal structures, a distance-preserving dimensionality reduction method is the most intuitive option. Multidimensional scaling (MDS) minimizes the difference between the input distances and the embedded distances, so it has previously been used for structural inference from Hi-C data (16–20). Though it is possible to use other dimensionality reduction methods to embed Hi-C data, the relationship between structural distances and physical distances is less clear with these methods.

**Figure 1.**
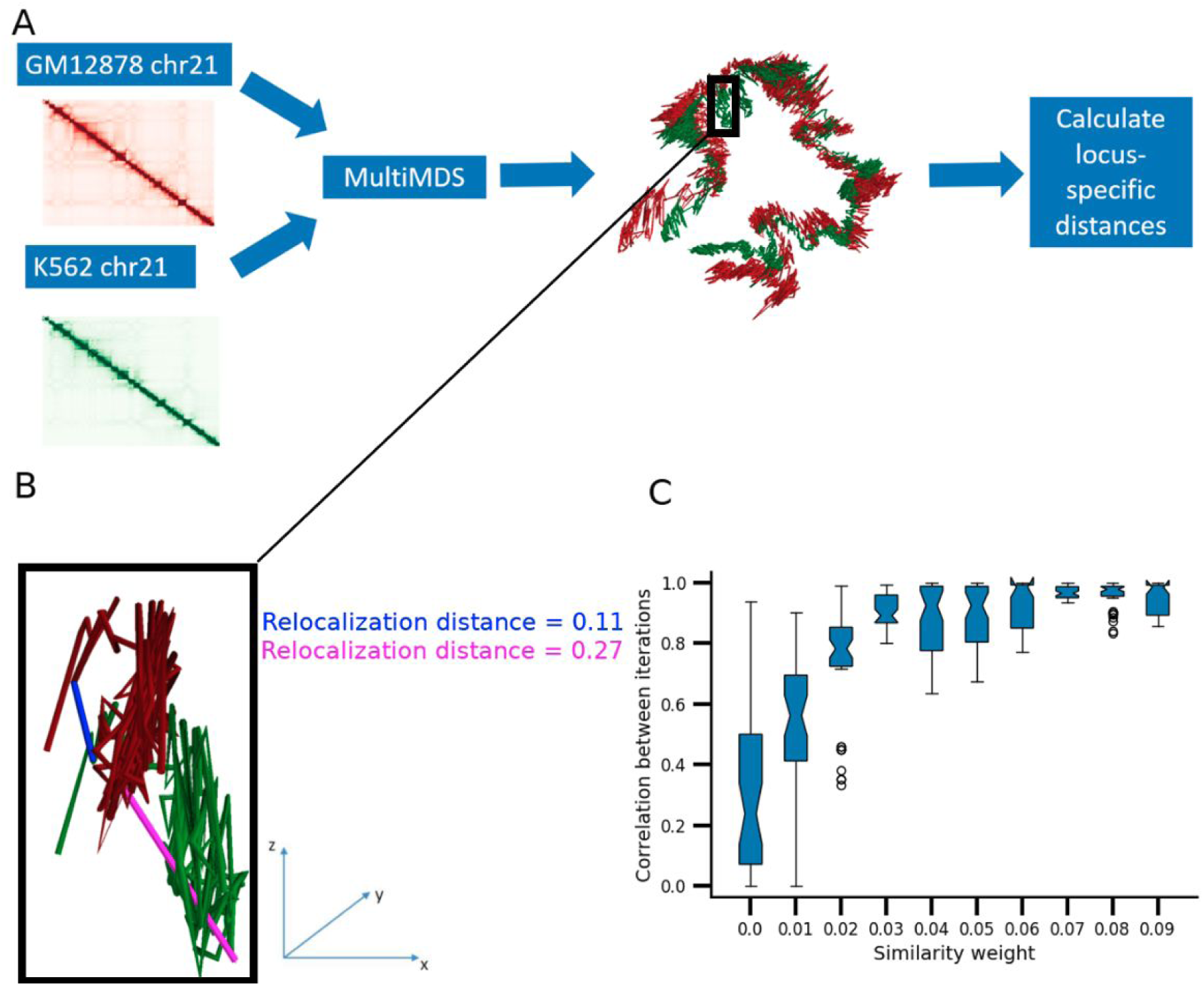
**A)** Example of MultiMDS applied to GM12878 and K562 chr21 datasets at 10kb resolution. **B)** Example of relocalization distance calculations in aligned GM12878 and K562 chr21 structures. Only chr21:22.26-23.26 Mb is shown. Relocalization distance is calculated for the loci 22.61-22.62 Mb (magenta) and 22.31-22.32 Mb (blue). (**C)** Pairwise correlations between multiple runs of MultiMDS applied to GM12878 and K562 chr21, measured across a range of difference penalties. Zero penalty represents independent inference and alignment.

It is possible to align independently estimated structures, for example (21). However, due to the inherent stochasticity of structural inference algorithms, independent structural inference followed by alignment overestimates the difference between datasets and results in irreproducible output. Though there are many Hi-C structural inference methods, ours is the first to jointly model two Hi-C datasets while sharing information between them.

To address the issue of stochasticity, MultiMDS performs MDS embedding and alignment simultaneously on both datasets, while minimizing the difference between embeddings multiplied by a similarity weight (Algorithm 1). The embedding difference term is easily incorporated into the MDS loss function (see methods), another advantage of using MDS over alternative dimensionality reduction methods. We tested the effects of various penalties on the alignment of GM12878 and K562 chr21. A similarity weight of zero is equivalent to independent MDS, so in this condition we performed Kabsch alignment after structural inference. We tested the effect of MultiMDS on reproducibility, measured as the correlation between pairwise MultiMDS output across multiple runs with the same input, at similarity weights between 0 and 0.1, to demonstrate that large similarity weights are not needed. Even a similarity weight of 0.05 suffices to improve the reproducibility of alignment to near perfect (Fig. 1C). On the other hand, even large similarity weights do not significantly worsen embedding accuracy relative to independent MDS, as the embedding error (measured as the root mean square difference between the input distance matrix and the embedding) for each dataset increases little even at a similarity weight of 0.5 (Supplementary Fig. 1), suggesting that there are multiple structures that fit the data similarly well. For another example, we tested mouse embryonic stem cell (mESC) and mouse hepatocyte chr19 with various difference penalties. A similarity weight of 0.02 suffices to improve reproducibility for this comparison, suggesting that these datasets are more similar (Supplementary Fig. 2). In general the optimal similarity weight depends on the datasets being compared and can be inferred by testing reproducibility at a variety of weights. MultiMDS provides the option to use the same partitioning algorithm as miniMDS to improve efficiency (22). Like miniMDS, MultiMDS is computationally efficient, with little increase in computational time relative to independent structure inference and alignment (Supplementary Fig. 3).

#### Algorithm 1 Joint multidimensional scaling

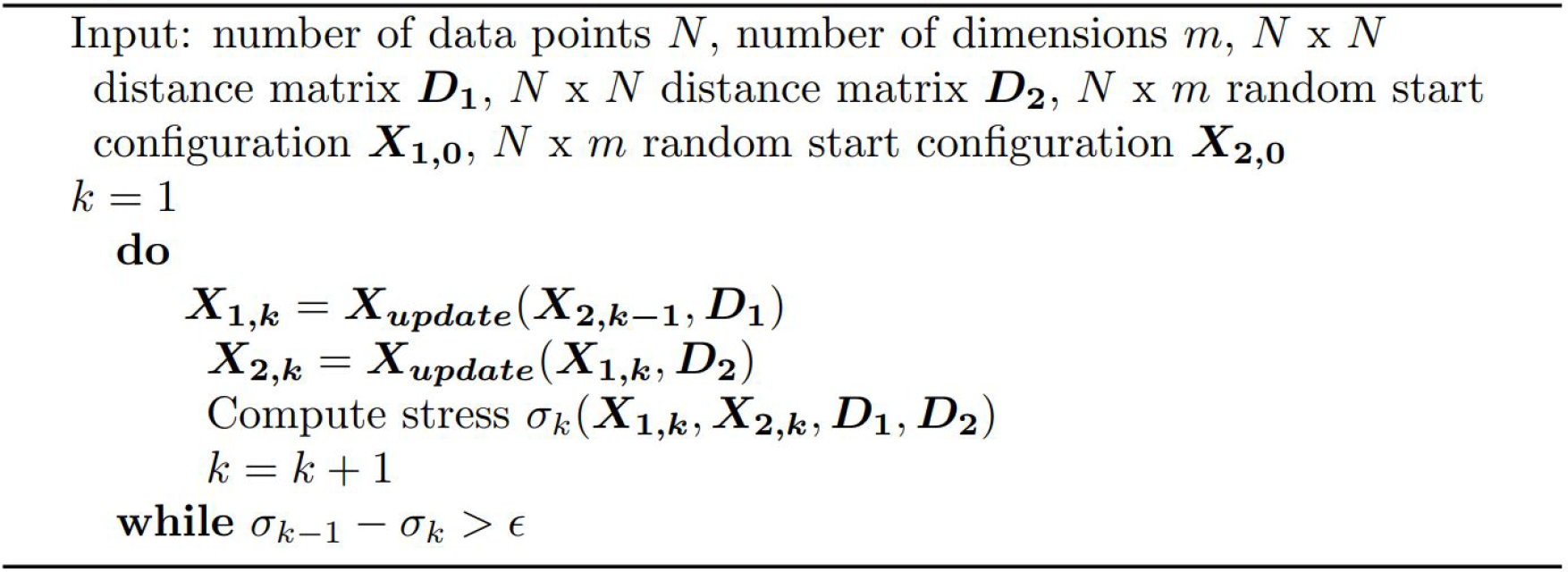

### MultiMDS analysis detects known galactose-dependent genome relocalizations in yeast

In order to demonstrate the abilities of MultiMDS to align chromosome structures and quantify locus-specific relocalization, we begin with comparisons of yeast Hi-C datasets. Chromatin structure reorganizes in yeast in response to changes in environment, but this has been difficult to systematically quantify because yeast do not have A/B compartmentalization to the extent that mammalian cells do. In mammalian cells long-range chromatin interactions are largely explained by compartment score, which is calculated as the first principal component of the interaction frequency correlation matrix (23). Compartment score correlates with the position of a locus along an axis between the active nuclear interior (A compartment) and the inactive lamina-associated domains (B compartment) (24) (Supplementary Fig. 4). Yeast lack a nuclear lamina (25) and so would not be expected to have A/B compartmentalization. As predicted, PC1 explained far less variance of the Hi-C correlation matrix in yeast compared to mouse and human (Supplementary Fig. 5A). Additionally, linear support vector regression (SVR) demonstrated that PC1 corresponds to a single physical axis in mouse and human, but not yeast, chromosomal structures (Supplementary Fig. 5B).

A previous study compared Hi-C data from yeast grown with glucose to yeast grown with galactose but was limited to measuring differential interaction frequency at loci of interest, which cannot identify loci that drive changes (26). Because the experiments were performed in hybrid yeast (*Saccharomyces cerevisiae* x *Saccharomyces uvarum*), it was possible to phase the data by homologs. The Has1-Tda1 locus was shown to pair with its homolog upon galactose induction.

MultiMDS comparison between glucose- and galactose-responsive Hi-C datasets appropriately detects relocalization of the Has1-Tda1 locus, but only for the *S. uvarum* homolog (Fig. 2A-B, Fig. 3A-B). The quantification suggests that only one locus drives pairing of the homologs, which could not have been determined by Hi-C loop calling. We also confirmed the expected relocalization of the Gal1-Gal7-Gal10 locus (Fig. 2C-D, Fig. 3C-D). MultiMDS showed that Gal3 and Gal4 also relocalize in the presence of galactose (Fig. 2E-H. Fig. 3E-H), though the relocalization of these genes had not been reported in the original study. The Gal1-Gal7-Gal10 relocalization was stronger for the *S. uvarum* homolog, whereas the Gal3 relocalization only occurred for the *S. cerevisiae* homolog. The Gal4 relocalization occurred for both homologs.

**Figure 2.**
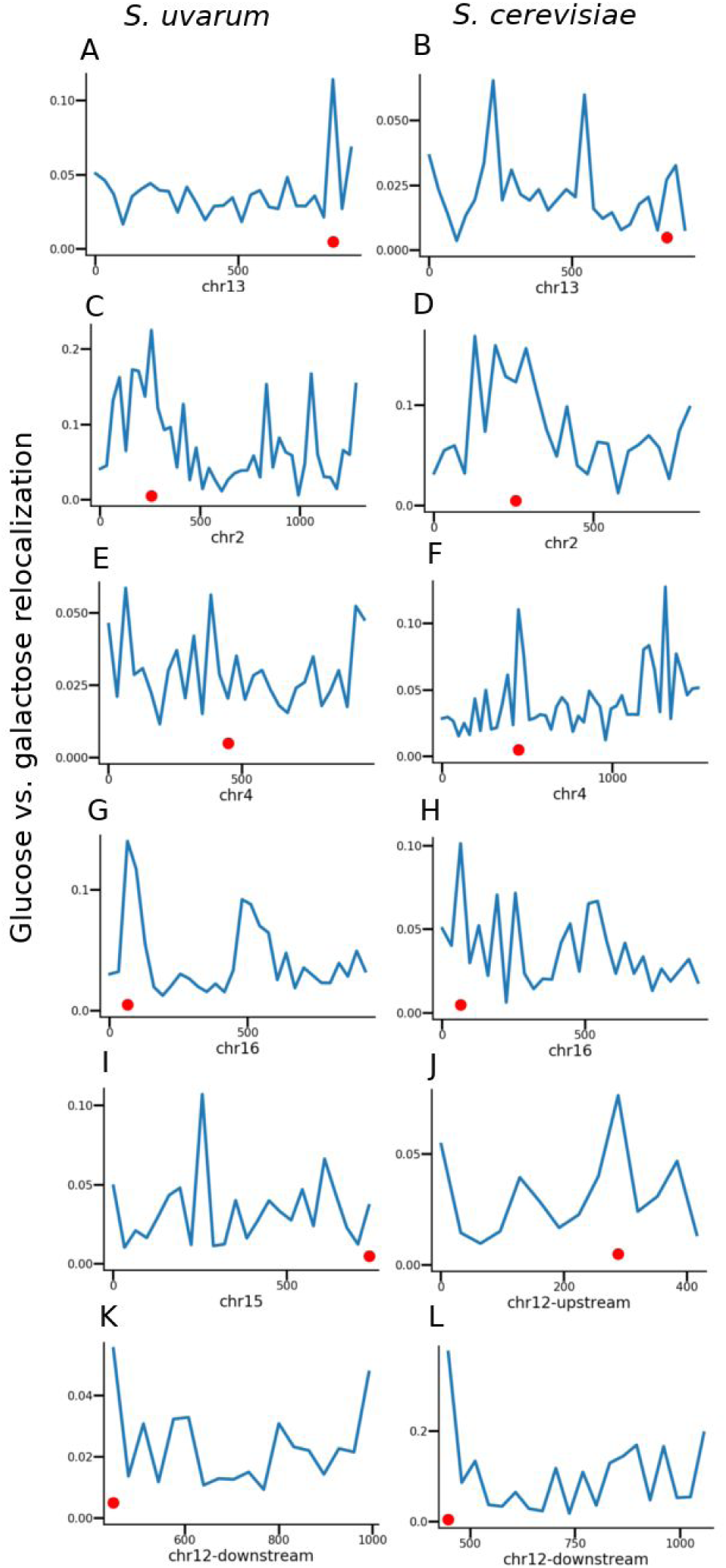
Relocalization of *S. cerevisiae* x *S. uvarum* hybrid yeast loci between glucose and galactose conditions, separated by homolog, for selected chromosomes. Genomic coordinates (kb) are shown on x axes. Loci of interest are highlighted with red dots.

**Figure 3.**
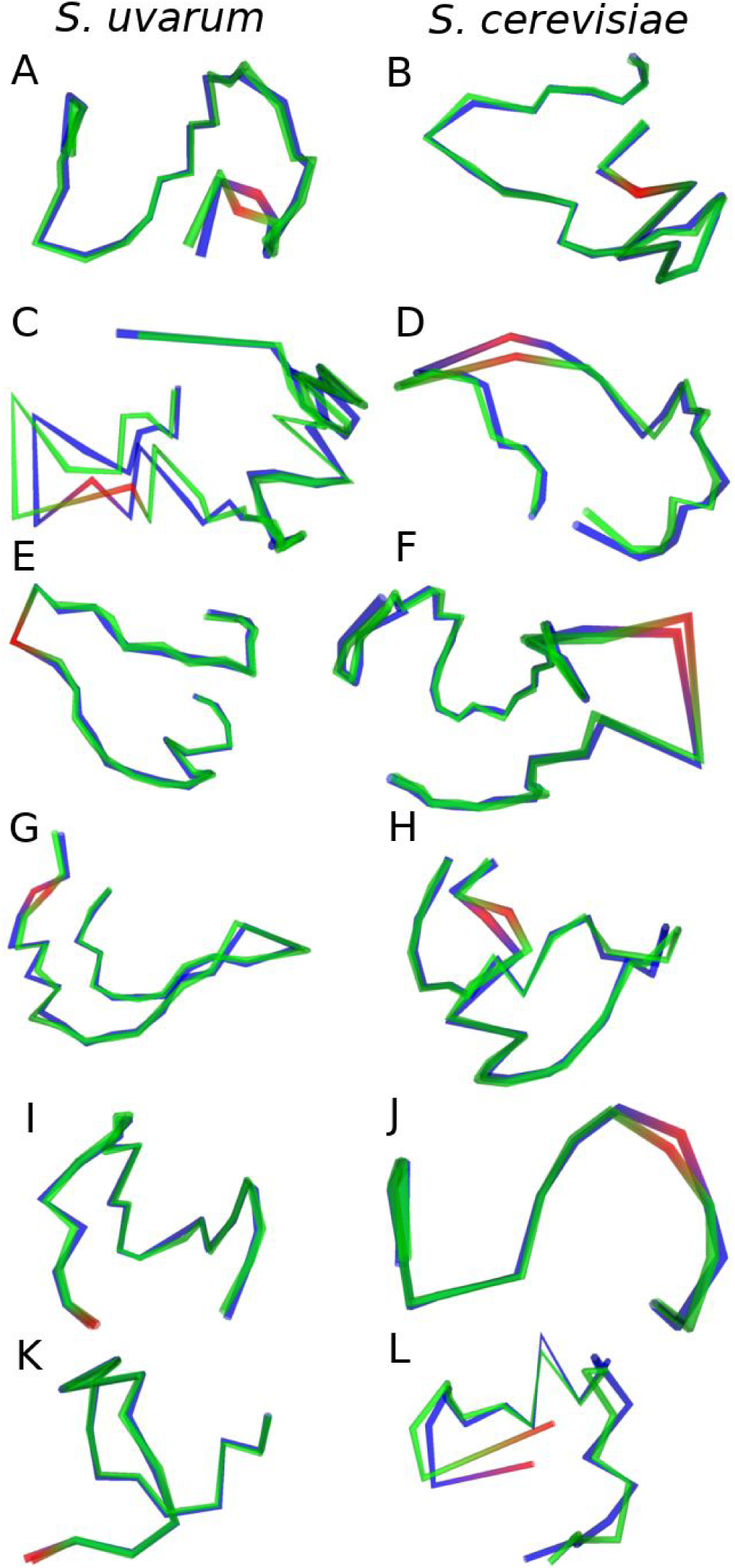
aligned structures for glucose (blue) and galactose (green) conditions. Loci of interest are highlighted in red (see fig. 2).

On chr12, we observed that the rDNA genes create a boundary between the upstream and downstream genomic regions. Because this made chr12 difficult to align, we aligned the two parts of the chromosome separately (Supplementary Fig. 6). In the upstream region, which excluded the rDNA genes, we observed relocalization of Gal2, with similar magnitude between homologs (Fig. 2I-J, Fig. 3I-J). In the downstream region, the *S. cerevisiae* rDNA genes displayed the strongest relocalization of any locus in the genome (Fig. 2K-L, Fig. 3K-L), which may be due to a change in conformation in the nucleolus. It has been previously shown that the yeast nucleolus changes conformation in different media, for example galactose or dextrose (27). Relocalization at these genes cannot be observed with independent structure inference and alignment, demonstrating the importance of MultiMDS’s joint structural inference (Supplementary Fig. 7). Though other examples of relocalized loci are seen in these datasets, the Gal genes and rDNA are among the highest peaks on their respective chromosomes.

Yeast genes, including Gal1-Gal7-Gal10, relocalize to the nuclear periphery upon activation, where they interact with the nuclear pore components, including Nup60 (26). Differential ChIP-seq enrichment of Nup60 in the presence of galactose relative to glucose had been qualitatively observed in Gal1-Gal7-Gal10, though not Has1-Tda1 (26). Using peak calling, we observed Nup60 peaks throughout the gene bodies of Gal1-7-10 (Supplementary Figs. 8A, 9A), Gal2 (Supplementary Figs. 8B, 9A), and Tda1 but not Has1 (Supplementary Figs. 8C, 9A). Differential enrichment was also found in the downstream portion of Gal3 (Supplementary Figs. 8D, 9A) and near the transcription start site of Gal4 (Supplementary Figs. 8E, 9A). As a negative control we found that Hxt1, a glucose transporter, lost Nup60 binding in the presence of galactose (Supplementary Figs. 8F, 9A). Tda1 and all relocalized Gal genes were upregulated, and Has1 and Hxt1 were downregulated (Supplementary Fig. 9B).

### Inter-compartment relocalizations dominate mammalian cell-specific differences in genome structure

Despite heterogeneity in chromosome conformation between individual cells, distinct patterns of compartmentalization can be observed between cell types, which correlate with differences in gene regulation (2,3). The detection of compartment changes in population Hi-C data suggests that localization relative to the nuclear periphery, and possibly other landmarks, could be detected using MultiMDS.

We first tested whether MultiMDS was able to identify the compartmentalization axis. Compartment scores are not used as input to MultiMDS, so this represents a somewhat independent validation. We performed MultiMDS on GM12878 and K562 datasets for each chromosome and performed linear SVR on the compartment scores for each 3D coordinate in the aligned structures. The SVR axis on average explains 87% of the variance in compartment scores in the GM12878 and K562 aligned structures, supporting the hypothesis that compartmentalization represents a single physical axis in the nucleus (Supplementary Fig. 10). It appears that the chromosomal structures in the alignment have similar positions relative to the nuclear periphery. The high SVR coefficients demonstrate that MultiMDS alignments are capturing consistent features of nuclear organization, rather than superficial similarities between the structures. When SVR is performed on unaligned structures, on average only 41% of the variance is explained by a single axis.

As expected, differences along the compartment axis correlate with compartment score differences. For example, the antiviral genes Mx1 and Mx2 have a weaker A compartment score in K562 relative to GM12878, associated with a loss of activity, and can be observed relocalizing along the compartment-associated axis in MultiMDS-aligned structures between these cell types (Supplementary Fig. 11).

Though it is clear that cell types differ in compartment score at certain loci, standard methods cannot systematically identify compartment-independent differences and thus cannot quantify the extent to which compartment differences explain global differences in Hi-C data. We used MultiMDS to calculate the fraction of relocalization that occurred along the compartment axis. To exclude the possibility that the physical lengths of each axis confound the results, we divided each fraction by the axis length. We found that the compartment axis is overrepresented for relocalization in pairwise comparisons of ENCODE cell types (GM12878, K562, KBM7, HUVEC, HMEC, and NHEK), even when normalized to axis length (p = 2.8 × 10^−149^) (Fig. 4A). Comparisons of mouse cell types (G1E, HPC-7, mESC, and hepatocyte) revealed a similar magnitude of compartment overrepresentation, indicating that the overrepresentation is not species-specific (p = 10^−41^) (Fig. 4B). Compartment axis differences are also overrepresented when comparing lymphoblastoid cell lines (LCLs) from different individuals (p = 3.4 × 10^−36^) (Fig. 4C). The overrepresentation of the compartment axis cannot be observed using independent structure inference and alignment (Supplementary Fig. 12). The relative overrepresentation of compartment axis relocalizations was consistent regardless of the total magnitude of relocalization, which varied significantly (Fig. 4E).

**Figure 4.**
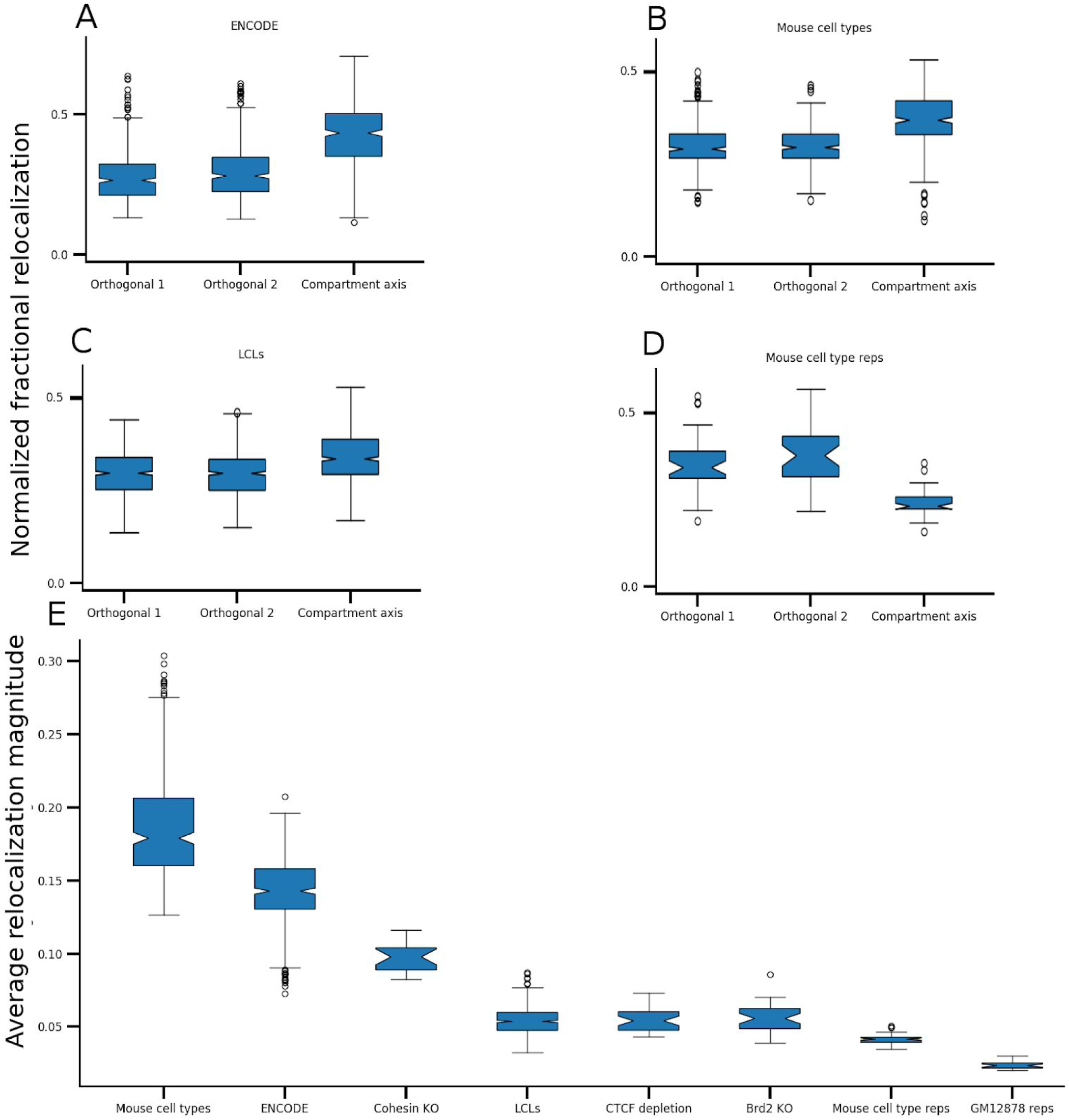
Fraction of total relocalization distance along each axis for all chromosomes for comparisons of **A)** ENCODE cell lines (GM12878, K562, KBM7, HUVEC, HMEC, and NHEK), **B)** mouse cell types (G1E, HPC-7, mESC, and hepatocyte), **C)** LCLs, and **D)** mouse cell type replicates. Total magnitude of difference along all axes is shown in **E**.

Given that compartmentalization is highly conserved between replicates (Supplementary fig. 13), we expect relocalizations between replicates to represent noise and not be enriched for compartment differences. Compartment axis overrepresentation was not seen when comparing GM12878 replicates (p = 0.08) (Supplementary fig. 14), and the compartment axis was in fact underrepresented in mouse cell type replicates (p = 3 × 10^−9^) (Fig. 4D). The underrepresentation may be due to the compartment axis being more constrained relative to the other axes, which may have more random variation.

Next we used MultiMDS to validate the relationship between various architectural proteins and compartmentalization. CTCF (28) and Brd2 (29) enforce TAD boundaries, and their loss does not affect compartmentalization. On the other hand, cohesin appears to oppose compartmentalization by forming TADs, and its loss causes a finer-grained compartmentalization to appear (30). We used MultiMDS to align Hi-C datasets from wild-type cells to Hi-C datasets resulting from depletions of CTCF, Brd2, and cohesin loading factor Nipbl. The relocalizations characterized by MultiMDS in comparisons between cohesin depletion and control Hi-C data are enriched along the compartment axis (p < 10^−4^) (Fig. 5A). Conversely, compartment axis relocalizations are not enriched in comparisons between the Brd2 depletion and control Hi-C data (p = 0.32) (Fig. 5B), and are slightly depleted in comparisons between the CTCF depletion and controls (p < 0.05) (Fig. 5C). The results of MultiMDS serve as validation of previous findings about the role of these proteins in the 3D genome.

**Figure 5.**
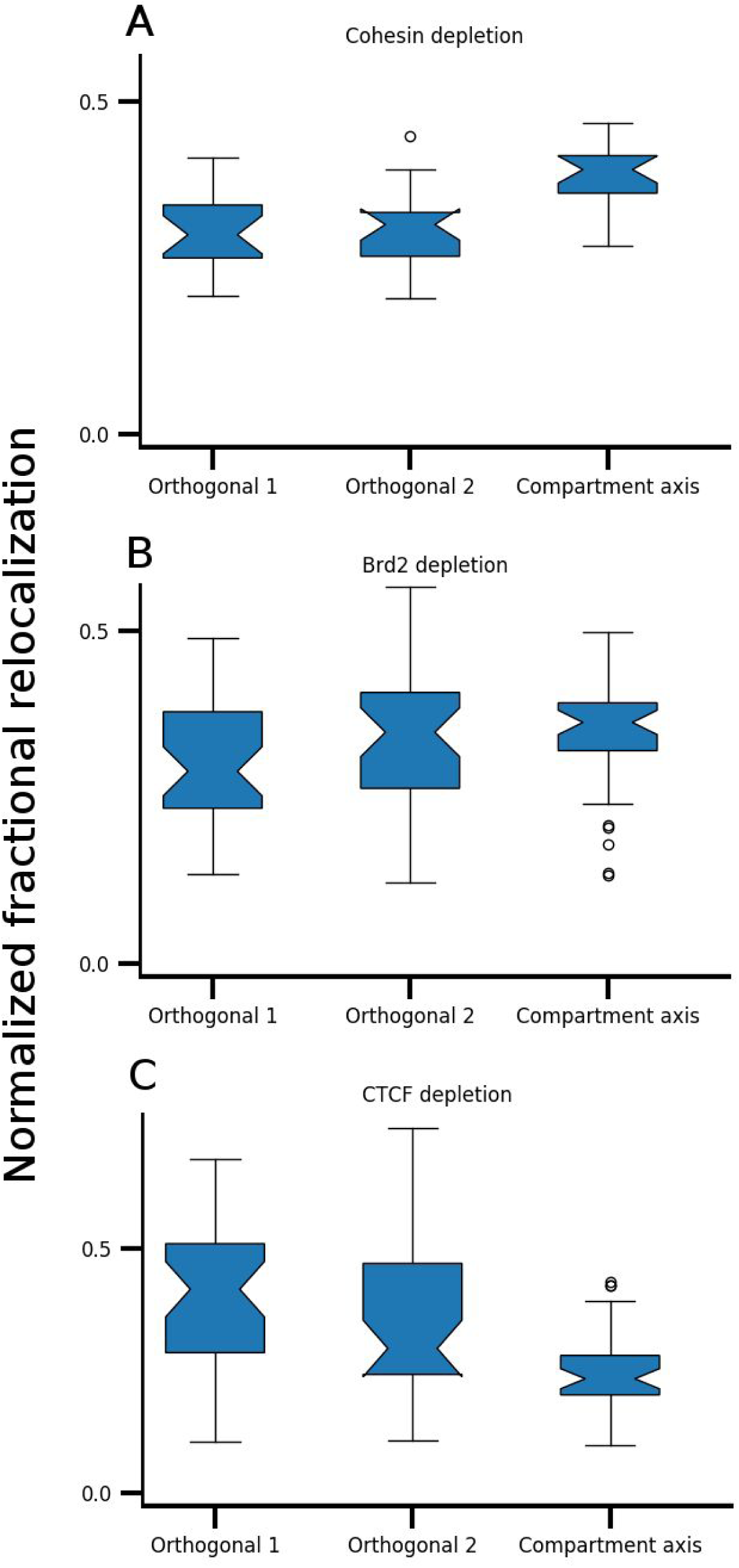
Fraction of total relocalization distance along each axis for all chromosomes upon depletion of architectural proteins: **A)** cohesin, **B)** Brd2, and **C)** CTCF.

### MultiMDS detects intra-compartment relocalizations across cell types

Despite the overrepresentation of relocalization along the compartment axis, some orthogonal relocalization also occurs. For example, most relocalization peaks overlap with differential compartment score peaks in comparisons between chr21 structures from IMR90 (embryonic lung fibroblast), HMEC (mammary epithelial), and HUVEC (umbilical vein endothelium) cell lines (Fig. 6A-C). However, some relocalization peaks do not overlap compartment difference peaks. In particular, a relocalization at chr21:47.4-47.5 Mb does not significantly differ in compartment score. This locus contains an intergenic region at chr21:47.75-47.5 Mb that displays different histone modifications across these cell lines (Supplementary Fig. 15A). In IMR90 the locus contains several accessible regions displaying H2A.Z, H3K4me1, H3K4me2, and H3K27ac ChIP-seq enrichment, DNase accessibility, and IDEAS enhancer states. HMEC cells also have active marks at this locus, though weaker and without H3K27ac, and has shorter regions of IDEAS enhancer states, as well as polycomb and heterochromatin states. HUVEC has low active ChIP-seq and DNase-seq signals at this locus, with some H3K27me3, and has more IDEAS polycomb and heterochromatin states and fewer enhancer states. As a negative control, we aligned the HUVEC data with K562 (chronic myelogenous leukemia), a cell line in which this locus has low active ChIP-seq and DNase-seq signal and high H3K27me3 signal and IDEAS polycomb and heterochromatin states. No relocalization was observed between K562 and HUVEC at this locus (Fig. 6D).

**Figure 6.**
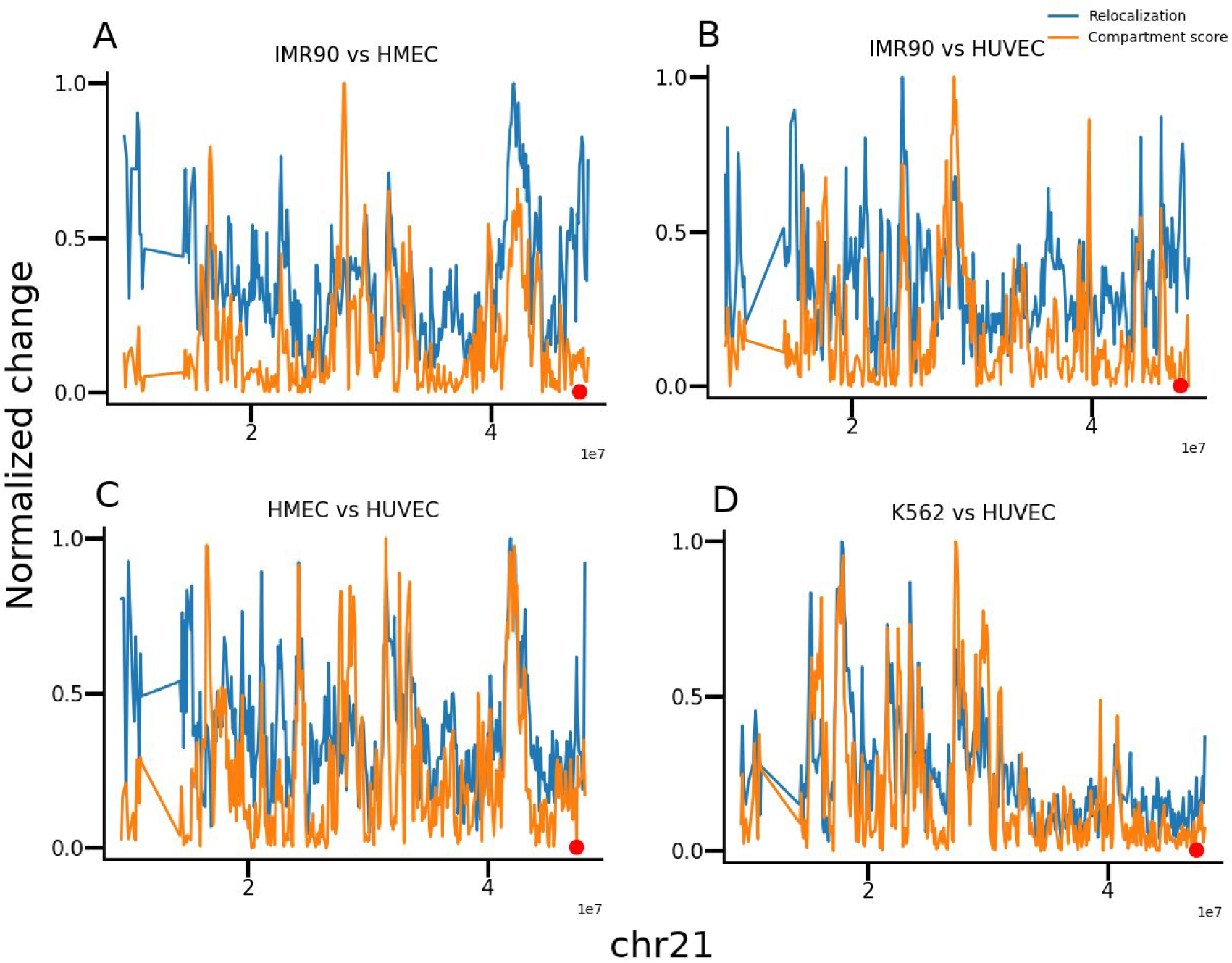
Relocalization and absolute compartment score differences (normalized to 1) between IMR90 and HMEC (A), HMEC and HUVEC (B), IMR90 and HUVEC (C), and K562 and HUVEC (D).

We then performed MultiMDS alignment at 25-kb resolution to visualize the conformation of the chr21:47.75-47.5 Mb locus. In IMR90, the locus appears to contact the chr21:46.9-47.0 Mb locus (Fig. 7A). In HMEC, the locus is closer to the chr21:46.9-47.0 Mb locus than in HUVEC and K562, but does not directly contact it (Fig. 7B-D).

**Figure 7.**
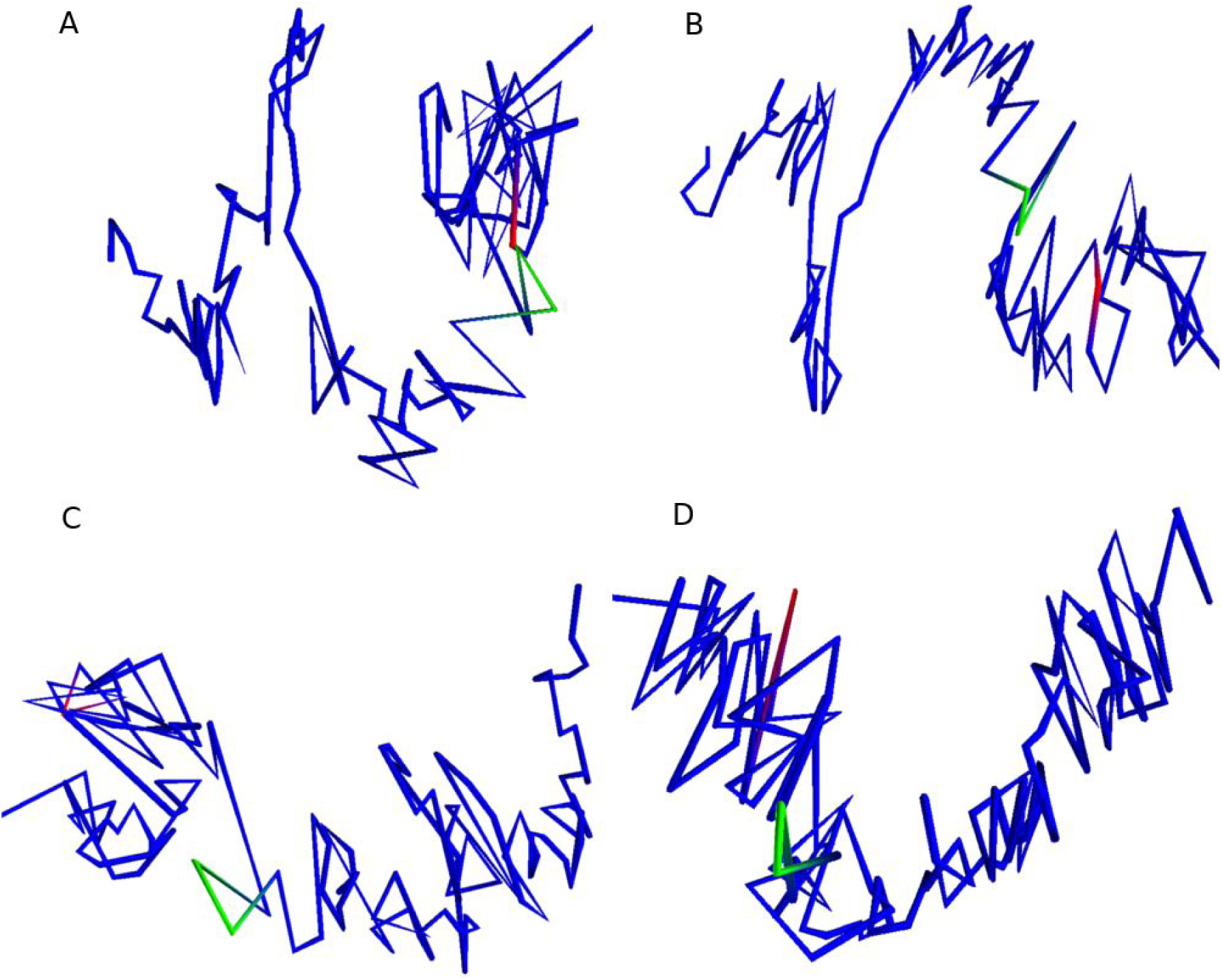
Structures for chr21:45.0-48.1 Mb in IMR90 (A), HMEC (B), HUVEC (C), and K562 (D). 47.4-47.5 Mb is highlighted in red, and chr21:46.9-47.0 Mb is highlighted in green.

Virtual 4C plots from the viewpoint of chr21:47.4-47.5 Mb reveal a strong peak at chr21:46.9-47.0 Mb in IMR90, the same region that the putative enhancer appears to contact in the 3D plots. This peak is present but weaker in HMEC (Fig. 8B) and not present in HUVEC and K562 (Fig. 8C-D). The chr21:46.9-47.0 Mb locus contains the COL18A1 gene, a component of the extracellular matrix, which has higher H3K36me3 signal in IMR90 relative to the other cell lines (Supplementary Fig. 15B).

**Figure 8.**
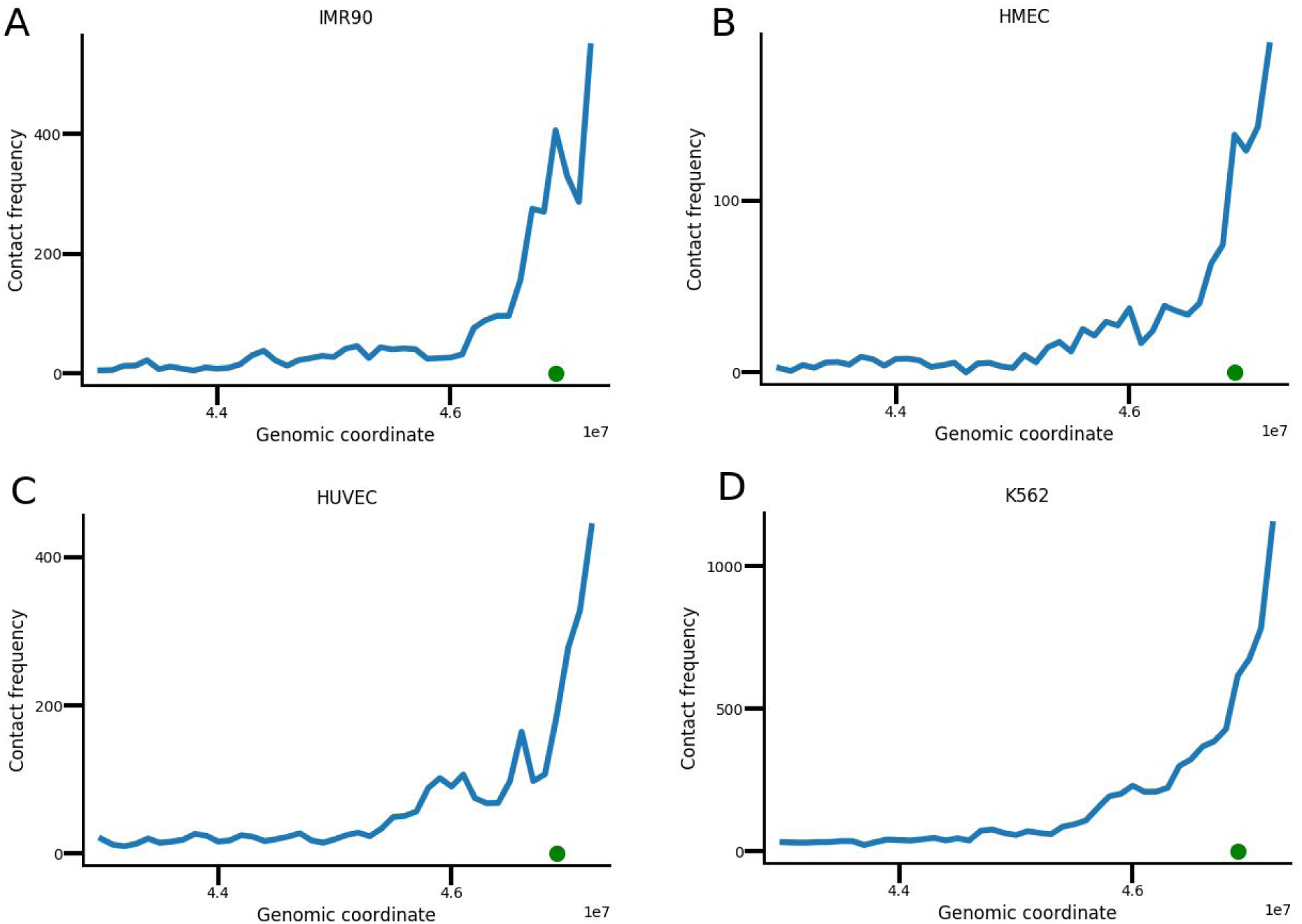
Virtual 4C plots with viewpoint at chr21:47.4-47.5 Mb for IMR90 (A), HMEC (B), HUVEC (C), and K562 (D). Green dot shows 46.9-47.0 Mb.

The differences in activity at the chr21:47.4-47.5 Mb locus cannot be predicted based on compartment scores alone. Though IMR90 has a slightly higher A compartment score at this locus compared to the other cell lines, HUVEC and K562 have higher compartment scores than HMEC, despite having less activity at this locus (Supplementary Table 1).

Next we performed a genome-wide quantification of relocalizations with minimal compartment score difference in GM12878 compared to K562. We called peaks in relocalization magnitude for 10-kb bins and, after filtering for mappability, identified 2562 peaks with absolute difference in compartment score of less than 0.2 (compartment scores range between −1 and 1) (Supplementary Fig. 16). Though some relocalization peaks do not overlap compartment differences, we noted that few compartment difference peaks occur without relocalization peaks, as would be expected. Because the B compartment has little biological signal, we focused on peaks that were within the A compartment in both cell types, which we term intra-A relocalization peaks. Relative to 35,636 mappability-filtered background loci in the A compartment, these peaks were enriched for H3K27ac, H3K4me1, H3K4me3, H3K9ac, H2A.Z, H3K4me2, and IDEAS enhancer states in GM12878, and depleted for these marks in K562 (Fig. 9). The peaks were enriched for the polycomb-associated marks H3K27me3 and EZH2 and IDEAS polycomb states in GM12878 and K562. H3K36me3 and IDEAS transcription states were depleted in both GM12878 and K562. These differences were not due to the peaks having different compartment scores or to differences in compartment score relative to the background (Supplementary Fig. 17). Most of these results cannot be observed using independent MDS (Supplementary Fig. 18).

**Figure 9.**
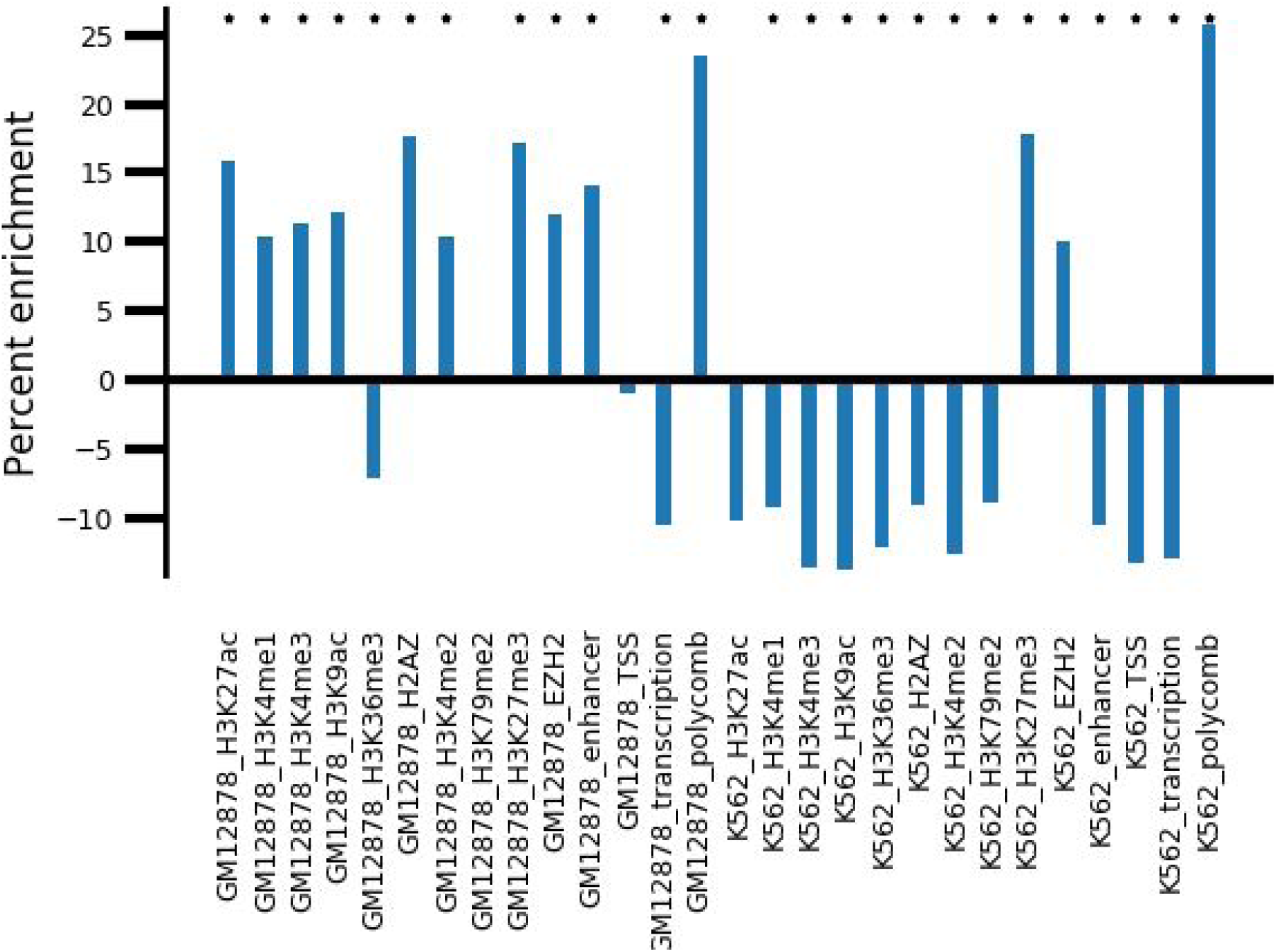
Enrichment of mean coverage of chromatin marks in intra-A relocalization peaks relative to background A compartment. Stars represent *p*<0.01.

Using hierarchical clustering of ChIP-seq coverage, we identified distinct subsets of intra-A relocalization peaks (Fig. 10). A large fraction of peaks are highly enriched for active marks. While some of these peaks are active in both GM12878 and K562, some lose active marks in K562 and gain H3K27me3. Other peaks have H3K27me3 in only one cell type but lack active marks in the other cell type.

**Figure 10.**
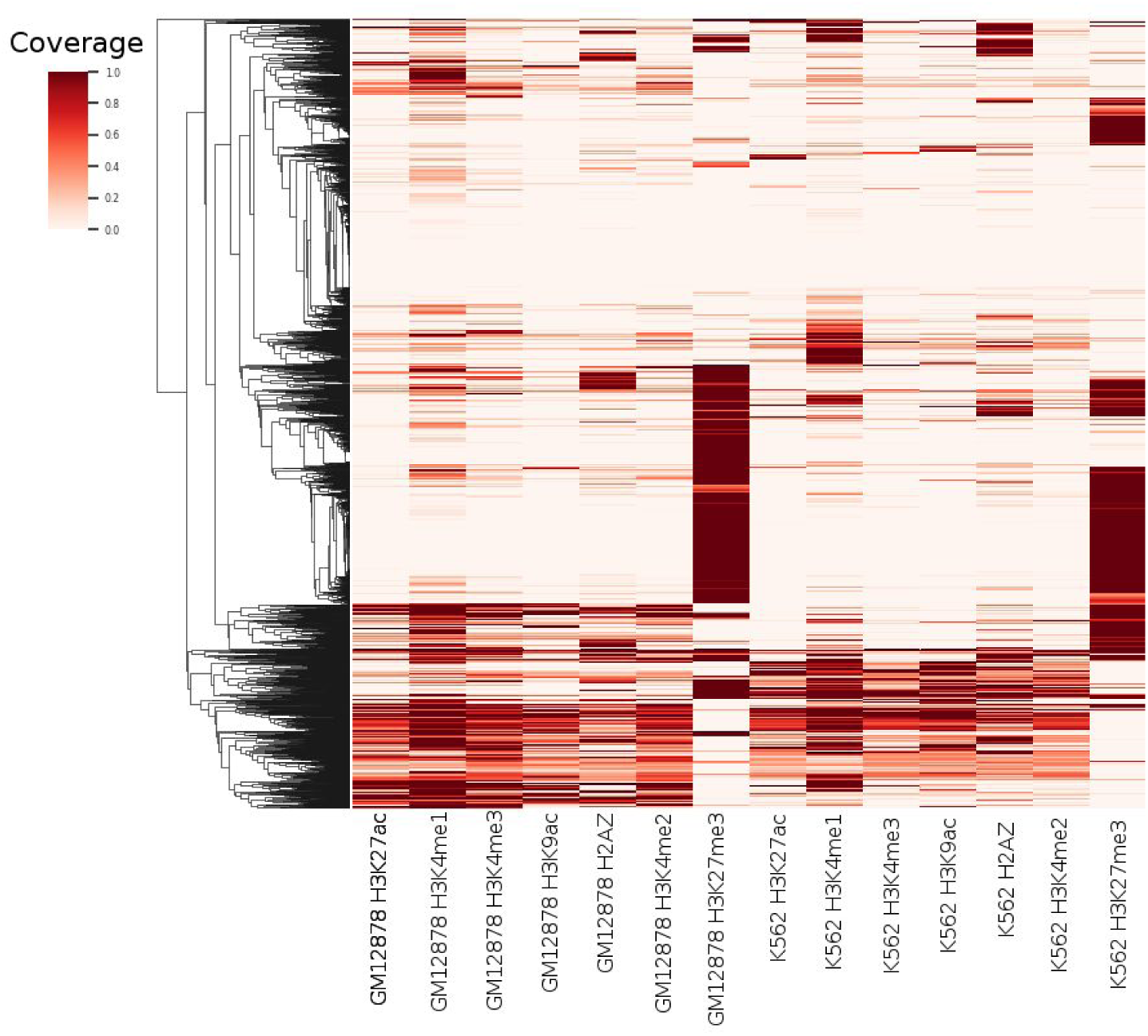
Hierarchical clustering of ChIP-seq peak coverage in intra-A relocalization peaks.

In summary, MultiMDS enables the detection of intra-compartment relocalizations, for example loci that change localization between two cell types but remain in the A compartment in both. Such intra-compartment relocalizations appear to be correlated with cell-specific changes in regulatory activity.

## DISCUSSION

MultiMDS is a computationally efficient tool for quantifying locus-specific differences between Hi-C datasets, which can be used even for Hi-C data that lacks compartment scores or TAD calls. It captures A/B compartment score differences better than alternative methods and quantifies the contribution of these differences to global differences in Hi-C data. At the same time, MultiMDS is able to go further than previous types of Hi-C analysis by identifying differences not based on differential compartmentalization, TADs, or looping. MultiMDS also differs from differential loop calling tools because it can identify the specific locus that drives loop formation. We demonstrated this ability by showing that a single homolog drives galactose-inducible homolog pairing at the Has1-Tda1 locus.

Because of the strong correlation between regulatory activity and compartmentalization, differences in compartment score have dominated previous comparisons of Hi-C data. To explore compartment-independent differences, we used Hi-C data from yeast, which lack compartmentalization. Examples of loci that change conformation upon galactose induction, such as Gal1-Gal7-Gal10 (26) and the nucleolus (27), had been previously identified, but comparisons had not been performed systematically. We confirmed these examples and also showed that Gal2, Gal3, and Gal4 relocalize in response to galactose. All the relocalized Gal genes are upregulated and gain Nup60 peaks, suggesting that the effect occurs via nuclear pore association. It had originally been reported that Has1-Tda1 does not gain Nup60 binding, based on visual inspection, but a quantitative analysis revealed a Nup60 peak at Tda1 upon galactose induction. The gain in nuclear pore association is consistent with the upregulation of Tda1 and its role in glucose metabolism (26).

Due to the strong effect of compartmentalization on long-range interactions, it is challenging to remove the effect of compartment score differences, which are strongly associated with differences in gene regulation, from other differences in genome structure. The contribution of compartment differences to overall structural differences was enriched in both mouse and human cell types and even in LCLs from different individuals. During the preparation of this manuscript, results were published showing that variation in chromosome conformation between LCLs is correlated with gene regulation (31). The overrepresentation of the compartment axis may be because differences along orthogonal axes are less functional and more stochastic and are thus less visible in aggregate Hi-C data. Consistent with this hypothesis, differences along the compartment axis are depleted in comparisons of mouse cell type replicate Hi-C datasets. As expected, a large fraction of changes after cohesin depletion are driven by compartment changes, specifically a gain in compartmentalization strength, while Brd2 depletion and CTCF depletion do not significantly affect compartmentalization.

Despite the enrichment of compartment axis relocalizations, we identified examples of loci that relocalized without significant compartment score differences. For example, a putative enhancer on chr21 relocalizes between cell types in which it appears active, poised, or polycomb repressed, suggesting that these states correspond to three distinct conformations. The presence of a cell-type-specific enhancer at this locus was consistent with the results of our genome-wide quantification of loci that relocalize within the A compartment between GM12878 and K562 with minimal compartment score difference. The relocalized loci were enriched for enhancers and polycomb in GM12878, suggesting active and poised enhancers, but were depleted for enhancers and enriched for polycomb in K562. The contrast between activity and repression may hint at global differences in regulation that occur between K562 and GM12878. On the other hand, the relocalized loci were depleted of active transcription in both cell types relative to the background A compartment. Given the strong relationship between compartment score differences and differential gene expression (2,3), the correlation with histone modifications further supports the hypothesis that the relocalizations we identify represent a compartment-independent regulatory mechanism.

The enrichment of enhancers at the relocalizations may be due to promoter-enhancer looping, as in the example of the enhancer on chr21. One possibility is that the enhancers are differentially associating with nuclear speckles. It has been shown that distance from nuclear speckles in the A compartment is independent of the nuclear lamina compartment axis and is correlated with superenhancers and H3K4me3, H3K9ac, and CTCF peaks (32). Thus the compartment-independent relocalizations we identified may represent differences in nuclear speckle association. Other relocalization peaks have cell-type-specific H3K27me3 without active marks, which may be associated with polycomb hubs that organize the 3D genome (33,34).

MultiMDS is a user-friendly tool that provides both visual and quantitative metrics of relocalization, as well as a built-in method for quantifying the contribution of the compartment axis to global differences, which could be used, for example, to determine the role of additional architectural proteins in compartmentalization. Though MultiMDS output cannot be interpreted as physical structures, MultiMDS is able to capture consistent structural features present throughout the population of cells. Our preliminary results showing the correlation of functional features with compartment-independent relocalizations hints at the existence of novel forms of nuclear organization, which can be further explored using MultiMDS. As more Hi-C datasets are produced, the number of possible comparisons will increase exponentially, improving our understanding of the relationship between 3D structure and function.

## METHODS

### Datasets

We used published Hi-C datasets for yeast grown in glucose and galactose media (Kim et al., 2017), ENCODE cell lines (35), lymphoblastoid cell lines (LCLs) (36) made available through the 4D Nucleome Project (37), wild-type and Brd2-knockout (KO) G1E cells (29), HPC-7 cells (38), wild-type and *Nipbl* conditional KO mouse hepatocytes (30), and wild-type and CTCF auxin-depleted mouse embryonic stem cells (mESCs) (28). Autosomes were used for all analyses. K562 chr9 and chr22 were removed due to their translocation.

Yeast RNA-seq read counts per gene and Nup60 ChIP-seq IP and input reads were from (Kim et al., 2017). Nup60 ChIP-seq data was aligned to the sacCer3 reference genome and broad peak calling was performed with MACS2. *Saccharomyces uvarum* gene annotations were from (Scannell et al., 2011). ChIP-seq data aligned to the hg19 reference genome was downloaded from www.encodeproject.org [access date?] (ENCODE Project Consortium, 2012). Replicated broad peak calls were used for relocalization enrichment analysis, and signal p-value was used for browser shots. IDEAS 20-state annotation based on Roadmap Epigenomics data was from (39).

### Algorithm

MultiMDS uses a novel joint multidimensional scaling (MDS) algorithm, which simultaneously embeds two distance matrices in a lower-dimensional space while minimizing the weighted sum of squared distances (SSD) between the embeddings. In addition to the weights for distance measurements used in standard MDS, which we refer to as distance weights, MultiMDS incorporates weights representing the expected similarity between the distance matrices, which we refer to as similarity weights. In our analysis, we used distance weights of 1 for all loci. We used equal similarity weights for all loci, equivalent to a single global parameter – referred to below as the penalty – which is selected empirically for each pairwise comparison (see below).

Assume we have *N* items observed under two conditions. Each condition has an *N* × *N* distance matrix. ***D*** The output of our algorithm is two *N* × *m* coordinate matrices ***X***_**1**_ and ***X***_**2**_, where *m* is the number of dimensions of the embedding (3 in our analyses).

The algorithm minimizes stress, which is calculated as

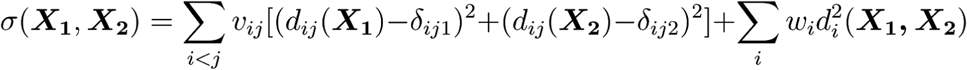

where *v*_*ij*_ is the distance weight for points *i* and *j, d*,_*ij*_(***X***_***a***_) is the Euclidean distance between points *i* and *j* in ***X***_***a***_, *δ*_*ija*_ is the dissimilarity between points *i* and *j* from distance matrix ***D***_***a***_,*w*_*i*_is the similarity weight for point *i*, and *d*_*i*_(***X***_1_, ***X***_2_) is the Euclidean distance between point *i* in distance matrix 1 and point *i* in distance matrix 2.

We seek to find a compact expression for SSD, 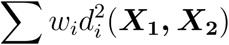, similar to (40). Consider one weighted squared distance, 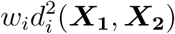, i.e. the distance for locus *i*.

Let ***x***_**1**_ ***a*** be column *a* of the coordinate matrix ***X***_**1**_, i.e. the *a*th dimension of the embedding. Let *e*_*i*_ be column *i* of the identity matrix ***I***.

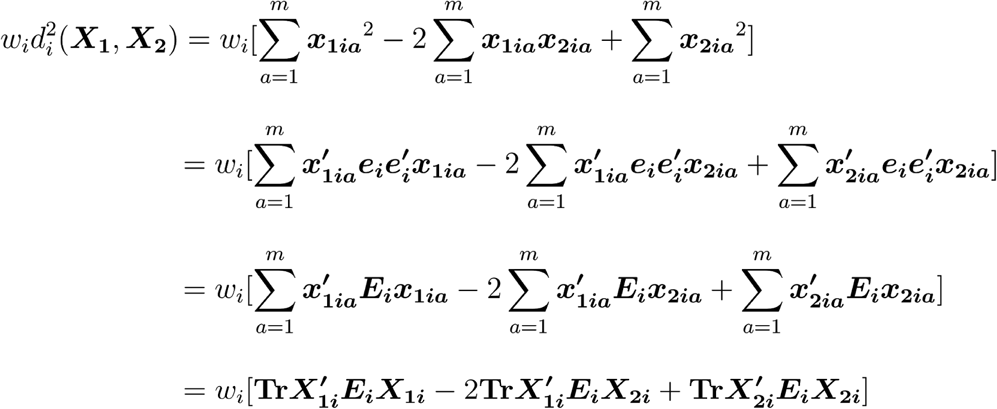

where ***E***_***i***_ is a matrix with *e*_*ii*_ = 1 and all other elements zero.

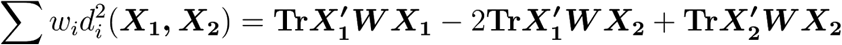

where ***W*** is a matrix with *w*_*i*_ = *w*_*i*_ and all other elements zero.

Combining with equation 8.27 in Borg & Groenen gives

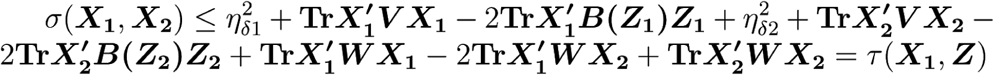

τ (***X***_**1**_,***Z***) achieves its minimum when ∇τ (***X***_**1**_,***Z***) = 0. Holding ***X***_**2**_ constant, we calculate the gradient and solve the system of linear equations as follows:

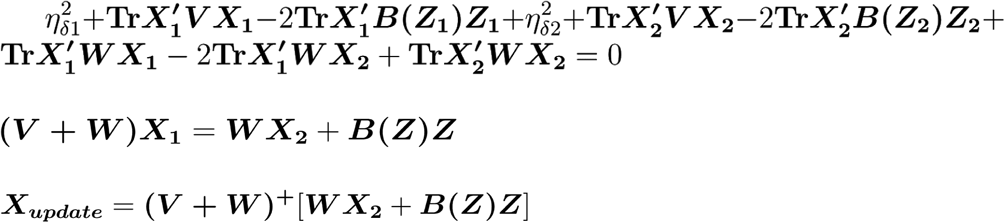

This is the update formula for ***X***_**1**_, where (***V***+***W***)^+^ is the Moore-Penrose inverse of ***V***+***W***. The update formula for ***X***_**2**_ is calculated similarly. The algorithm alternately updates ***X***_**1**_ and ***X***_**2**_ using the coordinates calculated at the previous step.

MultiMDS incorporates a small distance-decay prior to reduce noise. The corrected contact frequency is calculated as *c*_*corrected*_ = *c*_*observed*_ * (1 – *k*) + *c*_*expected*_ * *k*, where *c*_*observed*_ is the observed contact frequency, *c*_*expected*_ is the average contact frequency, for loci pairs with the same linear separation, and *k* is a weight parameter. *k* is set to 0.05 by default.

### Implementation

MultiMDS was implemented in python as a modification of miniMDS (22). The joint MDS algorithm was implemented using a modification of MDS from scikit-learn (41). By default, distances were calculated as (contact frequency)^-¼^. (Wang et al., 2015)

### Parameter selection

Similarity weights were selected by identifying the value at which increasing the similarity weight does not further increase reproducibility (see Fig. 1B).

### Comparison to independent structural inference and alignment

To compare MultiMDS with an alternate approach that aligns independently estimated genome structures, we first performed structural inference using miniMDS on each Hi-C dataset independently. The two structures were then aligned using the Kabsch algorithm (21), which minimizes the root mean square distance (RMSD) between two static structures.

### Compartment analysis

Compartment scores were calculated as described by (23). Briefly, a correlation matrix was calculated from the observed/expected Hi-C contact matrix. Compartment scores were defined as PC1 of the correlation matrix. PC1 accounts for most of the variance in the correlation matrix (up to 90% in high-coverage datasets). Scores were normalized to range between -1 and 1. The active compartment was defined as the compartment with greater enrichment for the IDEAS states 10_TssA, 14_TssWk, 8_TssAFlnk, 17_EnhGA, 6_EnhG, 4_Enh, 5_Tx, and 2_TxWk (39). The compartment axis for each chromosome was identified using linear support vector regression implemented in scikit-learn with default parameters (Pedregosa et al., 2011).

To prevent bias due to some axes being physically longer, the fraction of relocalization occuring along each axis was normalized by dividing by the axis length, which was calculated as the average distance of each coordinate from the centroid along that axis. Differences in normalized fractional relocalization were calculated using a two-sided independent-sample t-test.

### Relocalization peak analysis

Peaks in relocalization magnitude were called using a continuous wavelet transform (CWT) (Du et al., 2006), with peak widths ranging from 1 to 10. A CWT identifies peaks in noisy data based on their characteristic shape, without requiring smoothing.

Relocalization peaks with absolute compartment score differences of less than 0.2 were overlapped with ChIP-seq peaks and IDEAS states using bedtools to calculate coverage (Quinlan and Hall, 2010), and p-values were calculated using a two-sided independent sample t-test. The IDEAS states 17_EnhGA, 6_EnhG, 19_Enh/ReprPC, 18_Enh/Het, 4_Enh, and 11_EnhBiv were called enhancers, 16_TxRepr, 5_Tx, and 2_TxWk were called transcribed, and 13_ReprPC, 12_Het/ReprPC, and 1_ReprPCWk were called polycomb repressed.

### Differential ChIP-seq peaks

MACS2 (Zhang et al., 2008) was used to call broad peaks from Nup60 ChIP-seq data under glucose and galactose conditions separately. Glucose and galactose peaks were merged, and edgeR (Robinson et al., 2010) was run on the tag counts in each peak to identify differential enrichment between conditions. Because replicates were not available, the biological coefficient of variation was estimated as 0.1.

## DECLARATIONS

### Ethics approval and consent to participate

Not applicable

### Consent for publication

Not applicable

### Availability of data and materials

The scripts to reproduce figures and analysis are available at https://github.com/seqcode/multimds

### Authors’ contributions

LR implemented the algorithm and performed all analysis. LR and SM conceived the algorithm and wrote the manuscript. Both authors read and approved the final manuscript.

## Acknowledgements

The authors thank the members of the Center for Eukaryotic Gene Regulation at Penn State for helpful feedback and discussions.

## REFERENCES

1. Dixon JR, Selvaraj S, Yue F, Kim A, Li Y, Shen Y, et al. Topological domains in mammalian genomes identified by analysis of chromatin interactions. Nature. 2012 May 17;485(7398):376–80.

2. Dixon JR, Jung I, Selvaraj S, Shen Y, Antosiewicz-Bourget JE, Lee AY, et al. Chromatin architecture reorganization during stem cell differentiation. Nature. 2015 Feb 18;518(7539):331–6.

3. Lin YC, Benner C, Mansson R, Heinz S, Miyazaki K, Miyazaki M, et al. Global changes in the nuclear positioning of genes and intra- and interdomain genomic interactions that orchestrate B cell fate. Nat Immunol. 2012 Oct 14;13(12):1196–204.

4. Amano T, Sagai T, Tanabe H, Mizushina Y, Nakazawa H, Shiroishi T. Chromosomal Dynamics at the Shh Locus: Limb Bud-Specific Differential Regulation of Competence and Active Transcription. Dev Cell. 2009 Jan 20;16(1):47–57.

5. Ferraiuolo MA, Rousseau M, Miyamoto C, Shenker S, Wang XQD, Nadler M, et al. The three-dimensional architecture of Hox cluster silencing. Nucleic Acids Res. 2010 Nov 1;38(21):7472–84.

6. Rickman DS, Soong TD, Moss B, Mosquera JM, Dlabal J, Terry S, et al. Oncogene-mediated alterations in chromatin conformation. Proc Natl Acad Sci U S A. 2012 Jun 5;109(23):9083–8.

7. Sauria ME, Taylor J. QuASAR: Quality Assessment of Spatial Arrangement Reproducibility in Hi-C Data. bioRxiv. 2017 Nov 14;204438.

8. Ursu O, Boley N, Taranova M, Wang YXR, Yardimci GG, Stafford Noble W, et al. GenomeDISCO: a concordance score for chromosome conformation capture experiments using random walks on contact map graphs. Bioinformatics. 2018;2701–7.

9. Yan K-K, Yardimci GG, Yan C, Noble WS, Gerstein M. HiC-spector: a matrix library for spectral and reproducibility analysis of Hi-C contact maps. Bioinformatics. 2017 Jul 15;33(14):2199–201.

10. Yang T, Zhang F, Yardimci GG, Song F, Hardison RC, Noble WS, et al. HiCRep: assessing the reproducibility of Hi-C data using a stratum-adjusted correlation coefficient. Genome Res. 2017 Nov 1;27(11):1939–49.

11. Zufferey M, Tavernari D, Oricchio E, Ciriello G. Comparison of computational methods for the identification of topologically associating domains. Genome Biol. 2018 Dec 10;19(1):217.

12. Heinz S, Benner C, Spann N, Bertolino E, Lin YC, Laslo P, et al. Simple Combinations of Lineage-Determining Transcription Factors Prime cis-Regulatory Elements Required for Macrophage and B Cell Identities. Mol Cell. 2010 May 28;38(4):576–89.

13. Lun ATL, Smyth GK. diffHic: a Bioconductor package to detect differential genomic interactions in Hi-C data. BMC Bioinformatics. 2015 Aug 19;16:258.

14. Paulsen J, Sandve GK, Gundersen S, Lien TG, Trengereid K, Hovig E. HiBrowse: multi-purpose statistical analysis of genome-wide chromatin 3D organization. Bioinformatics. 2014 Jun 1;30(11):1620–2.

15. Stansfield JC, Cresswell KG, Vladimirov VI, Dozmorov MG. HiCcompare: an R-package for joint normalization and comparison of HI-C datasets. BMC Bioinformatics. 2018 Jul 31;19(1):279.

16. Duan Z, Andronescu M, Schutz K, McIlwain S, Kim YJ, Lee C, et al. A three-dimensional model of the yeast genome. Nature. 2010 May 20;465(7296):363–7.

17. Hirata Y, Oda A, Ohta K, Aihara K. Three-dimensional reconstruction of single-cell chromosome structure using recurrence plots. Sci Rep. 2016 Oct 11;6:34982.

18. Lesne A, Riposo J, Roger P, Cournac A, Mozziconacci J. 3D genome reconstruction from chromosomal contacts. Nat Methods. 2014 Nov;11(11):1141–3.

19. Morlot J-B, Mozziconacci J, Lesne A. Network concepts for analyzing 3D genome structure from chromosomal contact maps. EPJ Nonlinear Biomed Phys. 2016 May 5;4(1):2.

20. Shavit Y, Hamey FK, Lio P. FisHiCal: an R package for iterative FISH-based calibration of Hi-C data. Bioinformatics. 2014 Nov 1;30(21):3120–2.

21. Kabsch W. A solution for the best rotation to relate two sets of vectors. Acta Crystallogr A. 1976 Sep 1;32(5):922–3.

22. Rieber L, Mahony S. miniMDS: 3D structural inference from high-resolution Hi-C data. Bioinformatics. 2017 Jul 15;33(14):i261–6.

23. Lieberman-Aiden E, Berkum NL van, Williams L, Imakaev M, Ragoczy T, Telling A, et al. Comprehensive Mapping of Long-Range Interactions Reveals Folding Principles of the Human Genome. Science. 2009 Oct 9;326(5950):289–93.

24. Luperchio TR, Sauria ME, Wong X, Gaillard M-C, Tsang P, Pekrun K, et al. Chromosome Conformation Paints Reveal The Role Of Lamina Association In Genome Organization And Regulation. bioRxiv. 2017 Mar 30;122226.

25. Strambio-de-Castillia C, Blobel G, Rout MP. Isolation and characterization of nuclear envelopes from the yeast Saccharomyces. J Cell Biol. 1995 Oct 1;131(1):19–31.

26. Kim S, Liachko I, Brickner DG, Cook K, Noble WS, Brickner JH, et al. The dynamic three-dimensional organization of the diploid yeast genome. Ren B, editor. eLife. 2017 May 24;6:e23623.

27. Harris B, Bose T, Lee KK, Wang F, Lu S, Ross RT, et al. Cohesion promotes nucleolar structure and function. Mol Biol Cell. 2013 Dec 4;25(3):337–46.

28. Nora EP, Goloborodko A, Valton A-L, Gibcus JH, Uebersohn A, Abdennur N, et al. Targeted Degradation of CTCF Decouples Local Insulation of Chromosome Domains from Genomic Compartmentalization. Cell. 2017 May 18;169(5):930–944.e22.

29. Hsu SC, Gilgenast TG, Bartman CR, Edwards CR, Stonestrom AJ, Huang P, et al. The BET Protein BRD2 Cooperates with CTCF to Enforce Transcriptional and Architectural Boundaries. Mol Cell. 2017 Apr 6;66(1):102–116.e7.

30. Schwarzer W, Abdennur N, Goloborodko A, Pekowska A, Fudenberg G, Loe-Mie Y, et al. Two independent modes of chromatin organization revealed by cohesin removal. Nature. 2017 Nov;551(7678):51–6.

31. Gorkin DU, Qiu Y, Hu M, Fletez-Brant K, Liu T, Schmitt AD, et al. Common DNA sequence variation influences 3-dimensional conformation of the human genome. bioRxiv. 2019 Mar 30;592741.

32. Chen Y, Zhang Y, Wang Y, Zhang L, Brinkman EK, Adam SA, et al. Mapping 3D genome organization relative to nuclear compartments using TSA-Seq as a cytological ruler. J Cell Biol. 2018 Nov 5;217(11):4025–48.

33. Denholtz M, Bonora G, Chronis C, Splinter E, de Laat W, Ernst J, et al. Long-Range Chromatin Contacts in Embryonic Stem Cells Reveal a Role for Pluripotency Factors and Polycomb Proteins in Genome Organization. Cell Stem Cell. 2013 Nov 7;13(5):602–16.

34. Schoenfelder S, Sugar R, Dimond A, Javierre B-M, Armstrong H, Mifsud B, et al. Polycomb repressive complex PRC1 spatially constrains the mouse embryonic stem cell genome. Nat Genet. 2015 Oct;47(10):1179–86.

35. Rao SSP, Huntley MH, Durand NC, Stamenova EK, Bochkov ID, Robinson JT, et al. A 3D Map of the Human Genome at Kilobase Resolution Reveals Principles of Chromatin Looping. Cell. 2014 Dec 18;159(7):1665–80.

36. Chaisson MJP, Sanders AD, Zhao X, Malhotra A, Porubsky D, Rausch T, et al. Multi-platform discovery of haplotype-resolved structural variation in human genomes. bioRxiv. 2018 Jun 13;193144.

37. Dekker J, Belmont AS, Guttman M, Leshyk VO, Lis JT, Lomvardas S, et al. The 4D nucleome project. Nature. 2017 Sep;549(7671):219–26.

38. Wilson NK, Schoenfelder S, Hannah R, Castillo MS, Schütte J, Ladopoulos V, et al. Integrated genome-scale analysis of the transcriptional regulatory landscape in a blood stem/progenitor cell model. Blood. 2016 Jan 1;e12–23.

39. Zhang Y, An L, Yue F, Hardison RC. Jointly characterizing epigenetic dynamics across multiple human cell types. Nucleic Acids Res. 2016 Aug 19;44(14):6721–31.

40. Borg I, Groenen PJF. Modern Multidimensional Scaling: Theory and Applications. Springer Science & Business Media; 2005. 540 p.

41. Pedregosa F, Varoquaux G, Gramfort A, Michel V, Thirion B, Grisel O, et al. Scikit-learn: Machine Learning in Python. J Mach Learn Res. 2011 Oct;12:2825–30.

