## Supplemental information for "Joint inference and alignment of genome structures enables characterization of compartment-independent 3D relocalization across cell types"

**SUPPLEMENTARY FIGURES**

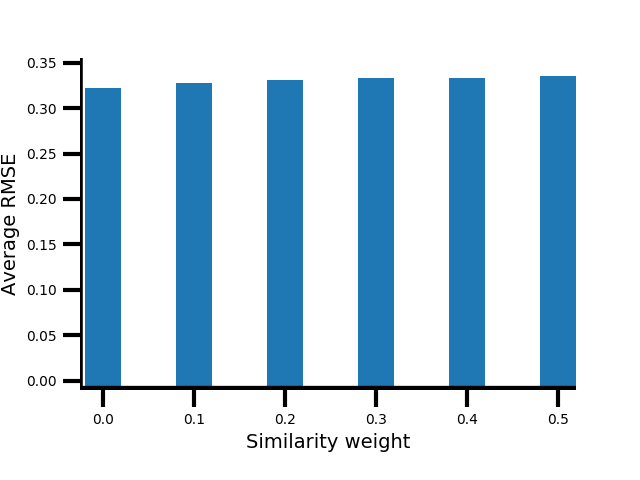

**Supplementary Figure 1.** Embedding error for MultiMDS run on GM12878 and K562 chr21, measured across a range of similarity weights.

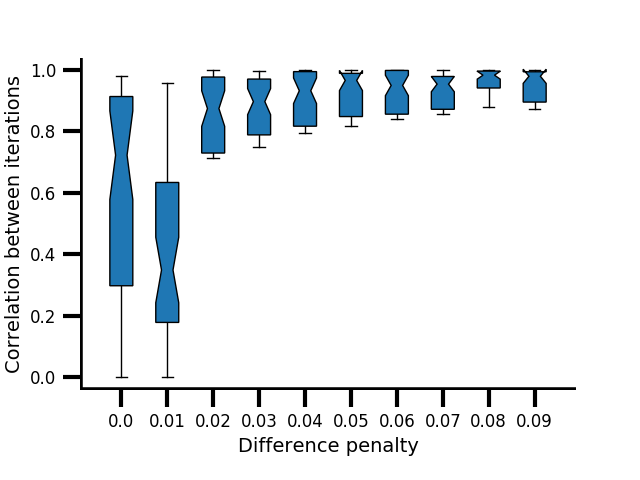

**Supplementary Figure 2.** Pairwise correlations between multiple runs of MultiMDS applied to mESC and mouse hepatocyte chr19, measured across a range of difference penalties. Zero penalty represents independent inference and alignment.

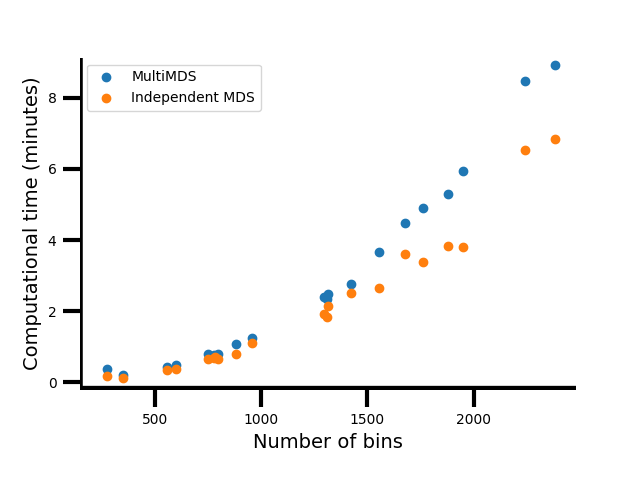

**Supplementary Figure 3.** Computational time for datasets of various sizes, measured for MultiMDS and for independent MDS structural inference followed by alignment.

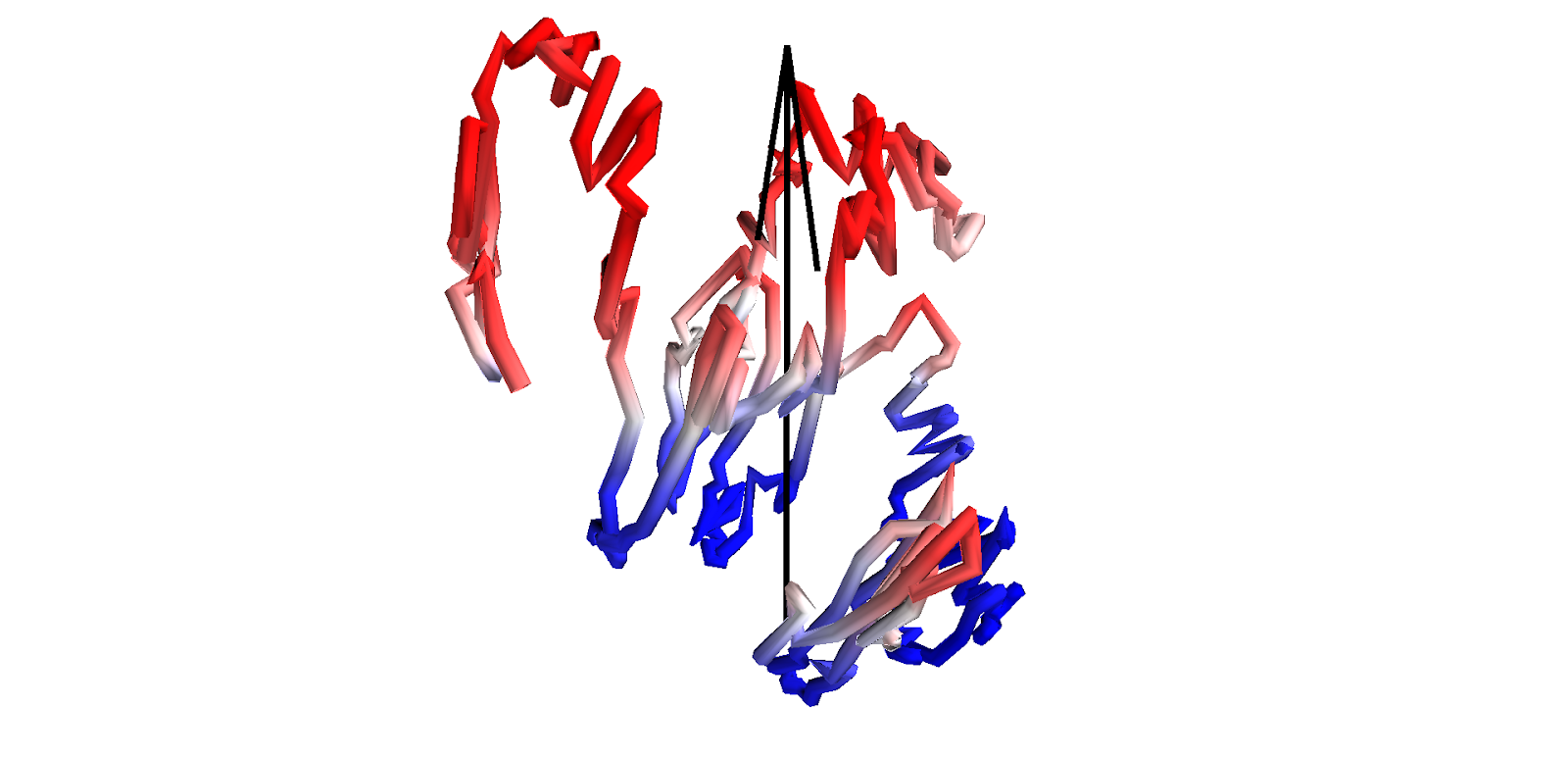

**Supplementary Figure 4.** Compartment score plotted on GM12878 chr21 at 100kb resolution. Positive scores (corresponding to A compartment) are plotted in reds, and negative scores (corresponding to B compartment) are plotted in blues. The structure has been rotated so that the SVR axis (arrow) aligns with the z axis. Pericentromeric regions have been removed.

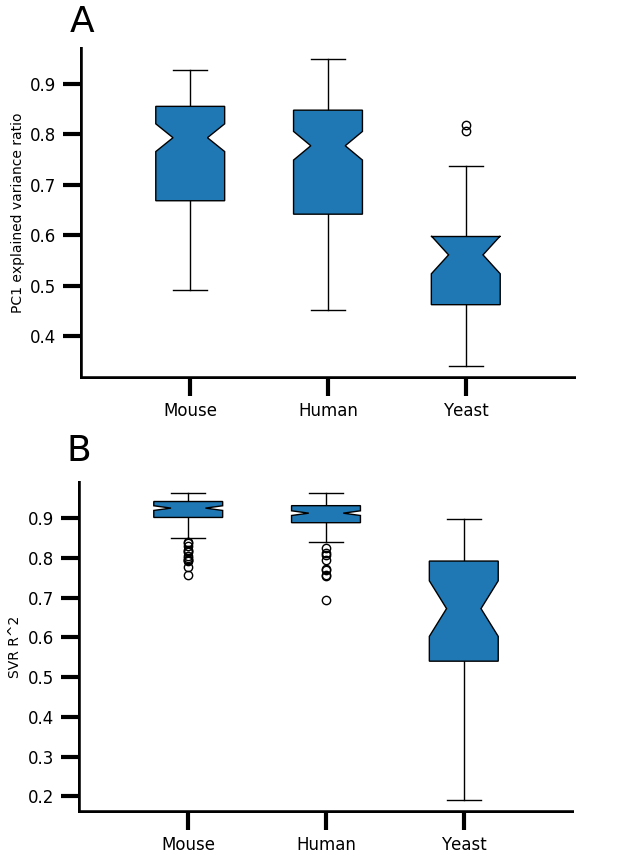

**Supplementary Figure 5.** A) Variance explained by PC1 of Hi-C correlation matrix by species. B) Linear SVR coefficient of determination of compartment scores predicted by 3D coordinates by species.

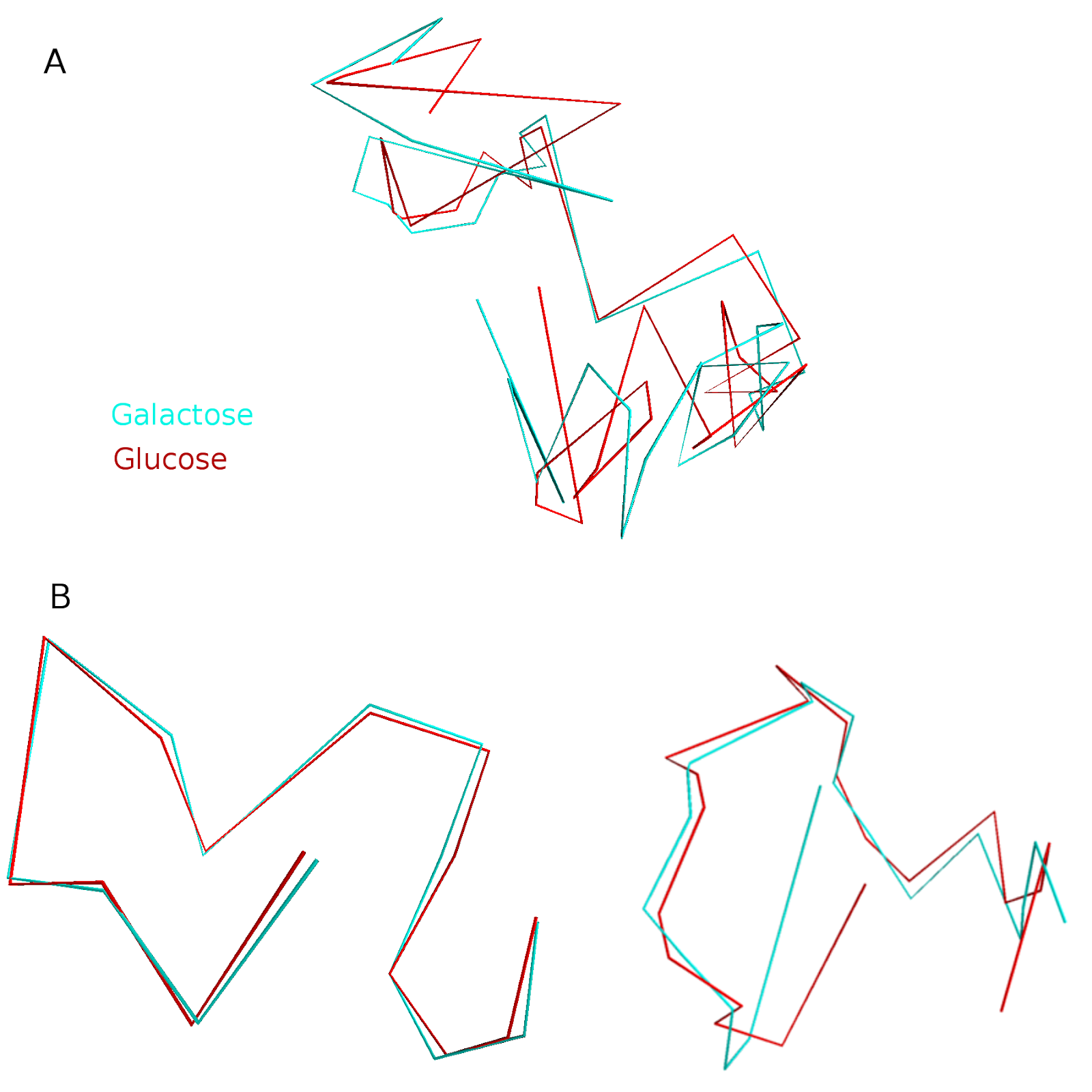

**Supplementary Figure 6.** chr12 *S. cerevisiae* glucose (red) and galactose (cyan) alignment for **A)** full structures and **B)** split structures.

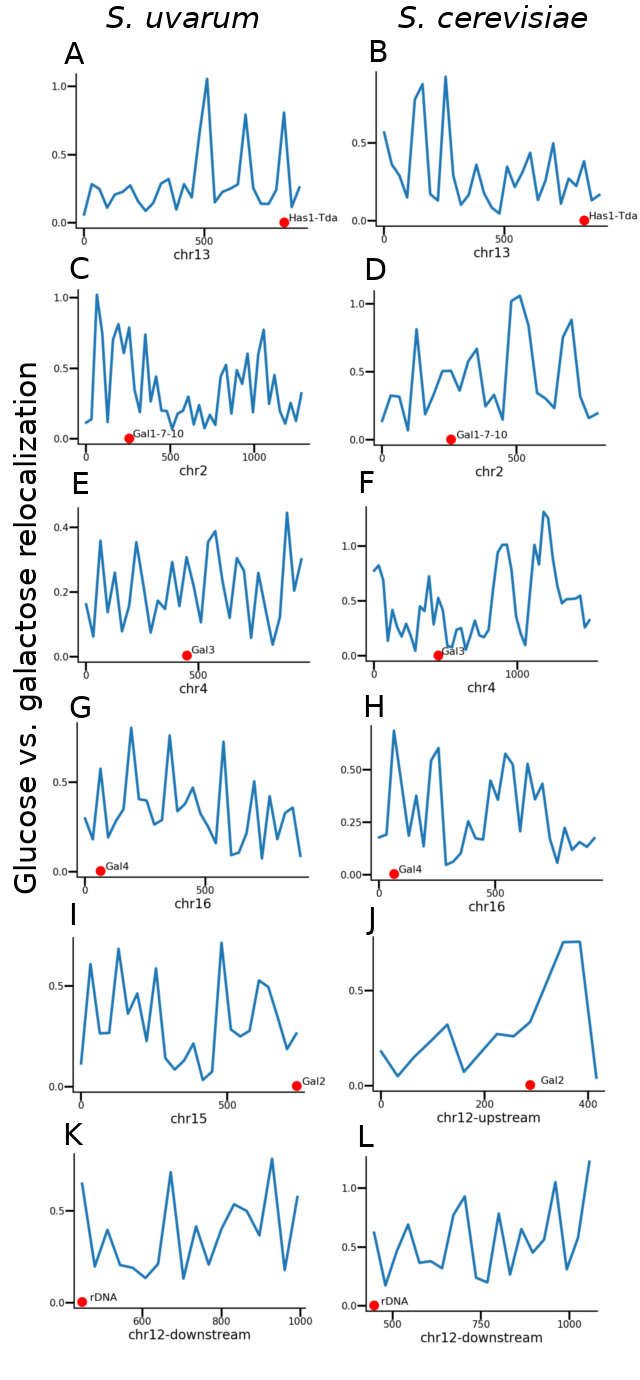

**Supplementary Figure 7.** Locus-specific relocalization calculated from independent structural inference and alignment. Genomic coordinates (kb) are shown on x axes. Loci of interest are highlighted with red dots.

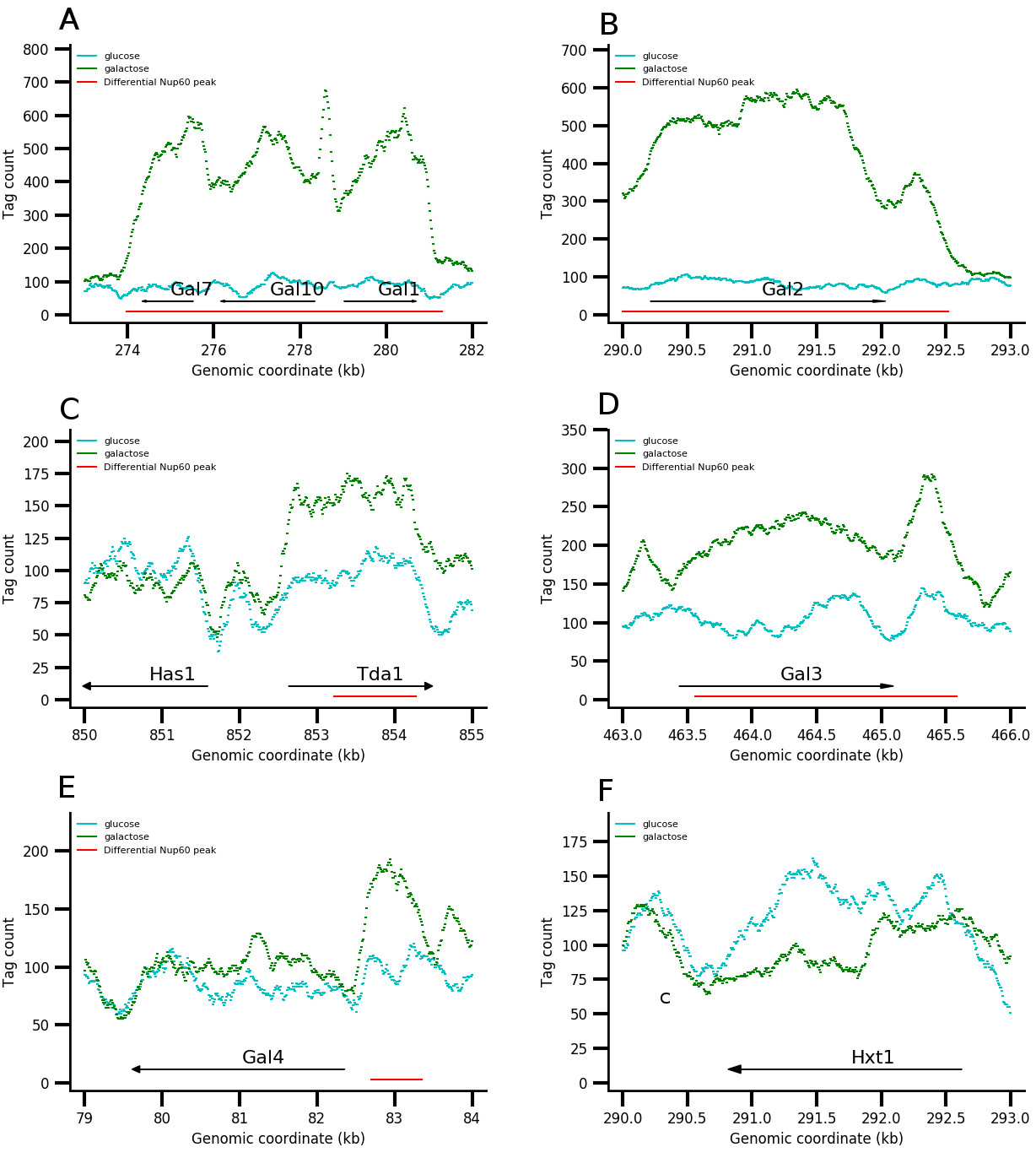

**Supplementary Figure 8.** Nup60 ChIP-seq tag counts in glucose and galactose conditions at **A)** Gal1-Gal7-Gal10, **B)** Gal2, **C)** Has1-Tda1, **D)** Gal3, **E)** Gal4, and **F)** Hxt1.

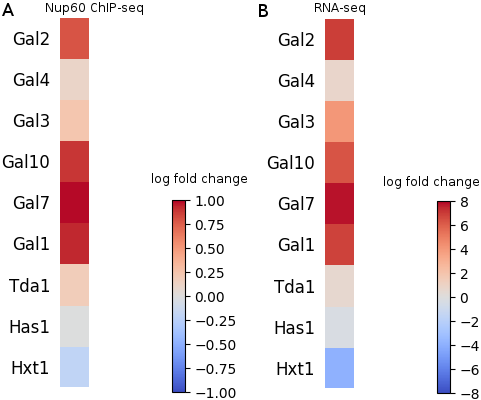

**Supplementary Figure 9.** Log fold change of **A)** Nup60 ChIP-seq tag counts and **B)** RNA-seq counts at selected genes in galactose relative to glucose.

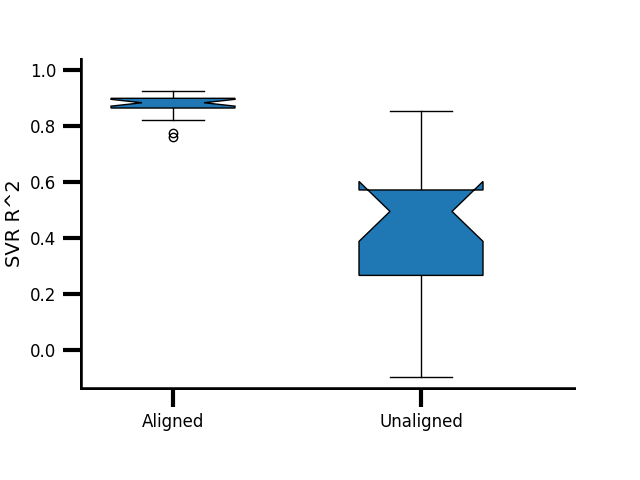

**Supplementary Figure 10.** SVR coefficient of determination of compartment scores predicted by 3D coordinates for aligned and unaligned GM12878 and K562 structures

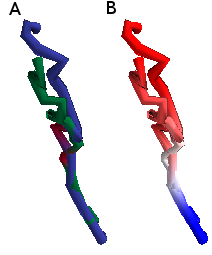

**Supplementary Figure 11.** chr21:41900000-43900000 aligned to the compartment axis (see Supplementary Fig. 7). **A)** Blue: GM12878. Green: K562: Red: Mx1/Mx2. **B)** Compartment scores. Positive scores are plotted in reds, and negative scores are plotted in blues.

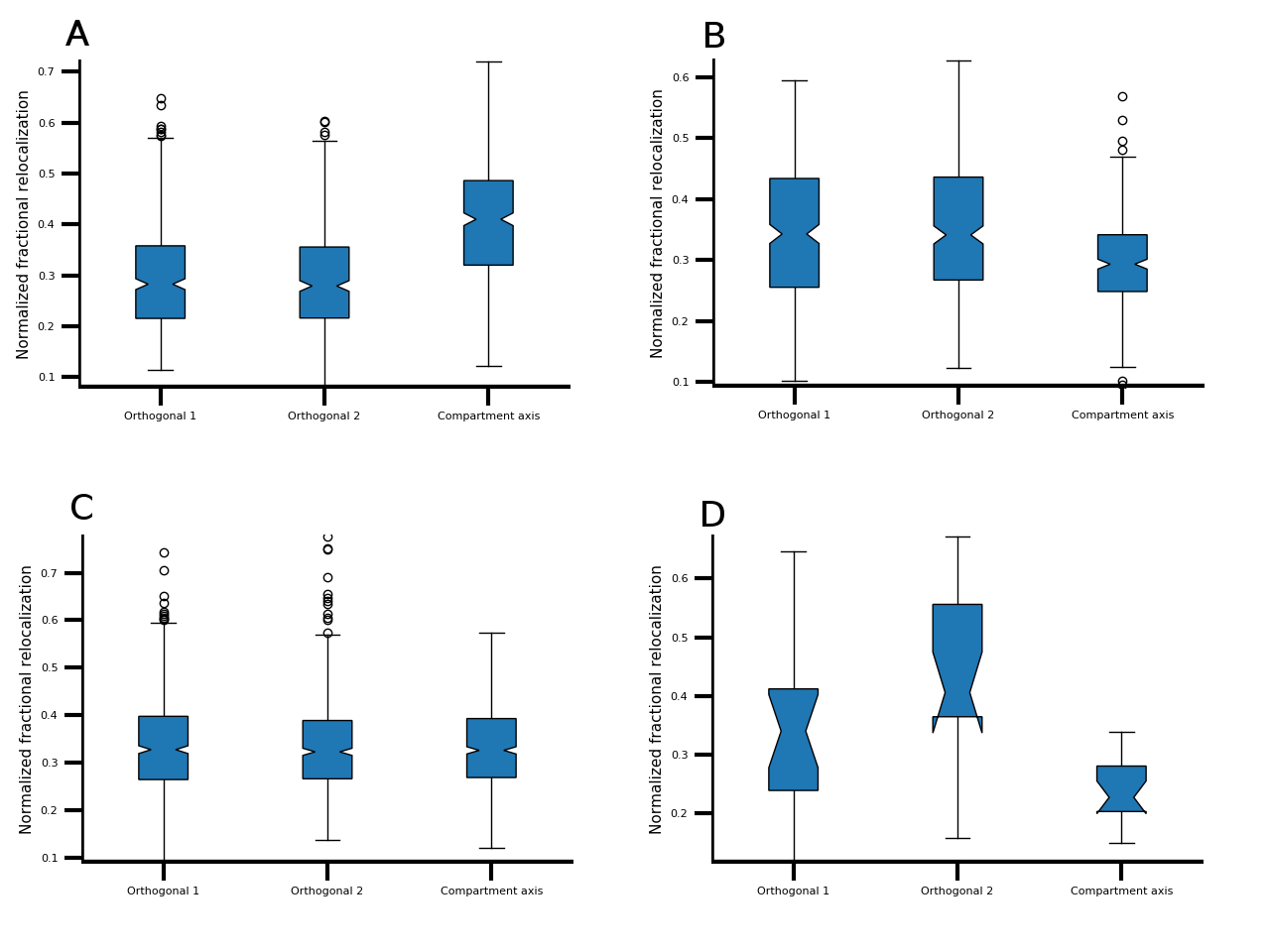

**Supplementary Figure 12.** Fraction of total relocalization distance along each axis for all chromosomes, calculated from independent structural inference and alignment, for **A)** ENCODE cell lines, **B)** mouse cell types, **C)** LCLs, and **D)** cohesin KO.

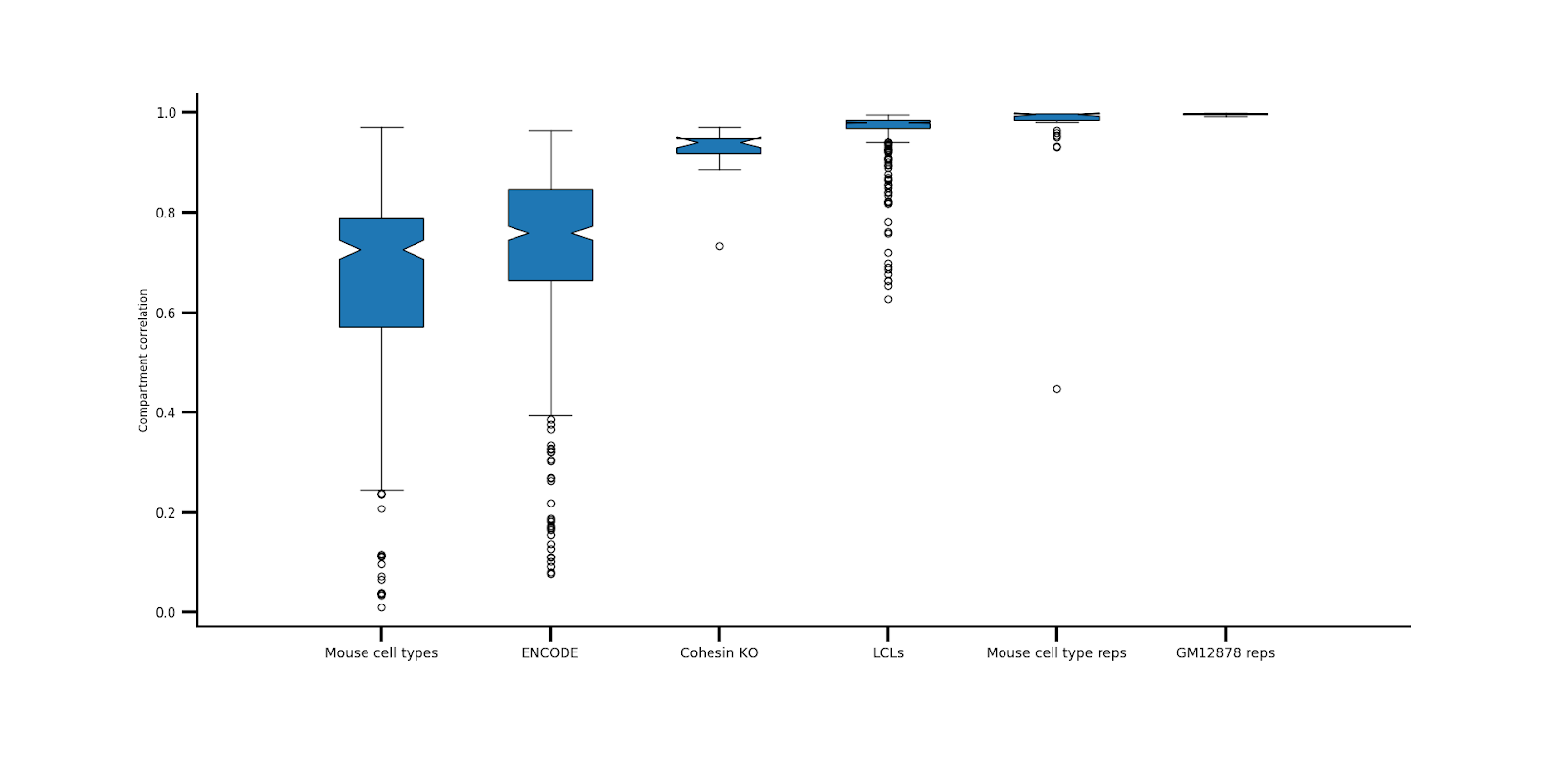

**Supplementary figure 13.** Compartment score correlations between each pair of datasets, separated by data type.

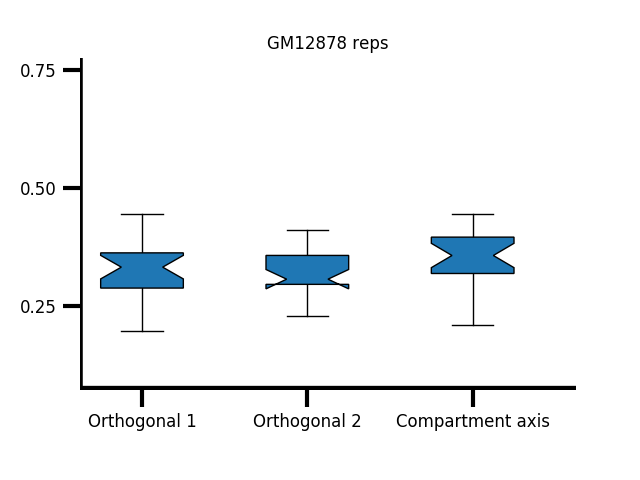

**Supplementary figure 14.** Fraction of total relocalization distance along each axis for all chromosomes for GM12878 replicates.

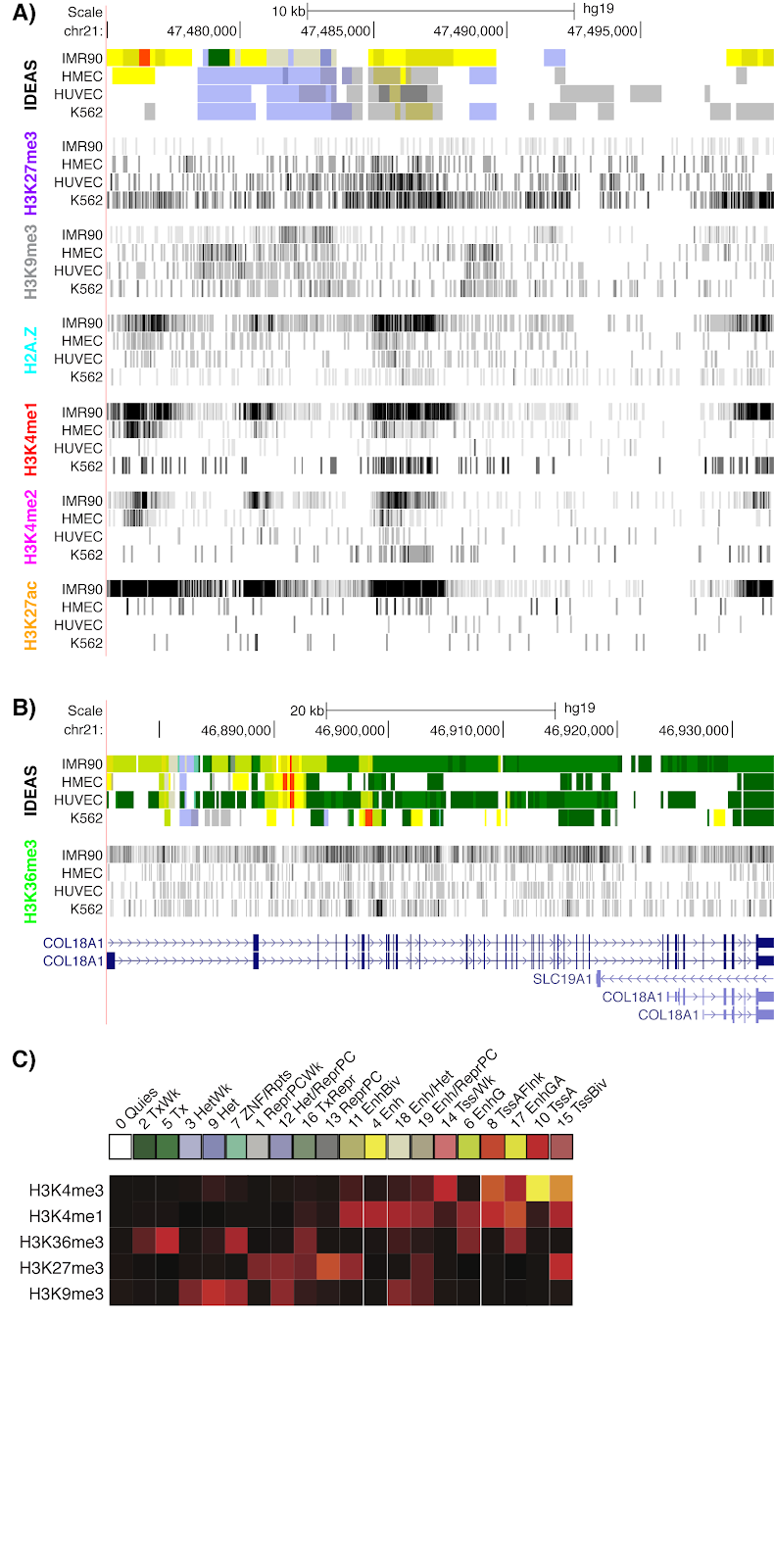

**Supplementary Figure 15.** A) ChIP-seq data for chr21:47.475-47.5 Mb. B) H3K36me3 ChIP-seq data for COL18A1. C) Emission probabilities for IDEAS states.

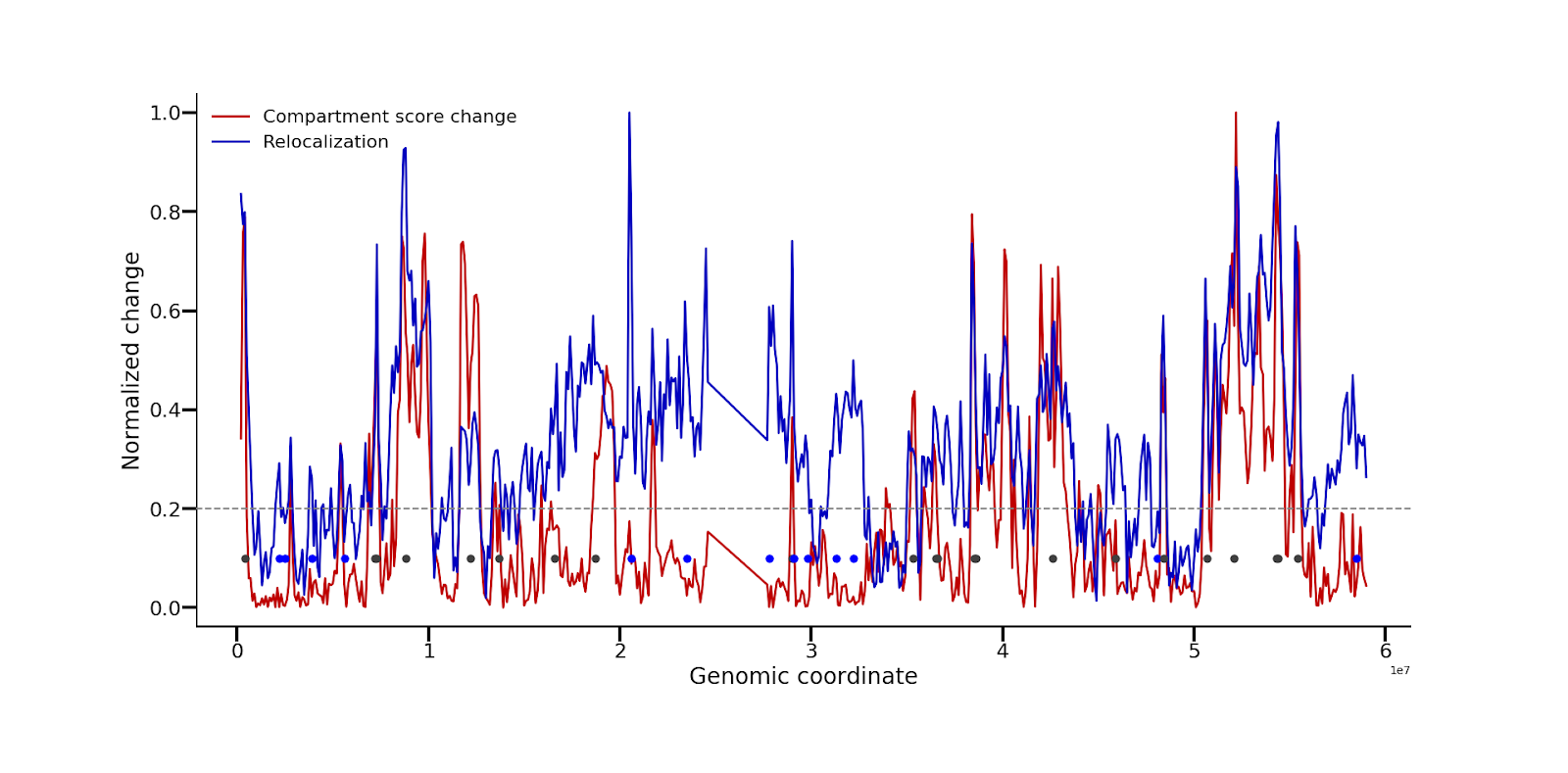

**Supplementary Figure 16.** Relocalization and absolute compartment score differences between GM12878 and K562 chr19, normalized to 1. For ease of visualization, 100kb resolution is shown. Gray dashed line is the threshold for compartment-independent relocalizations. Gray dots are compartment-dependent relocalizations, and blue dots are compartment-independent relocalizations.

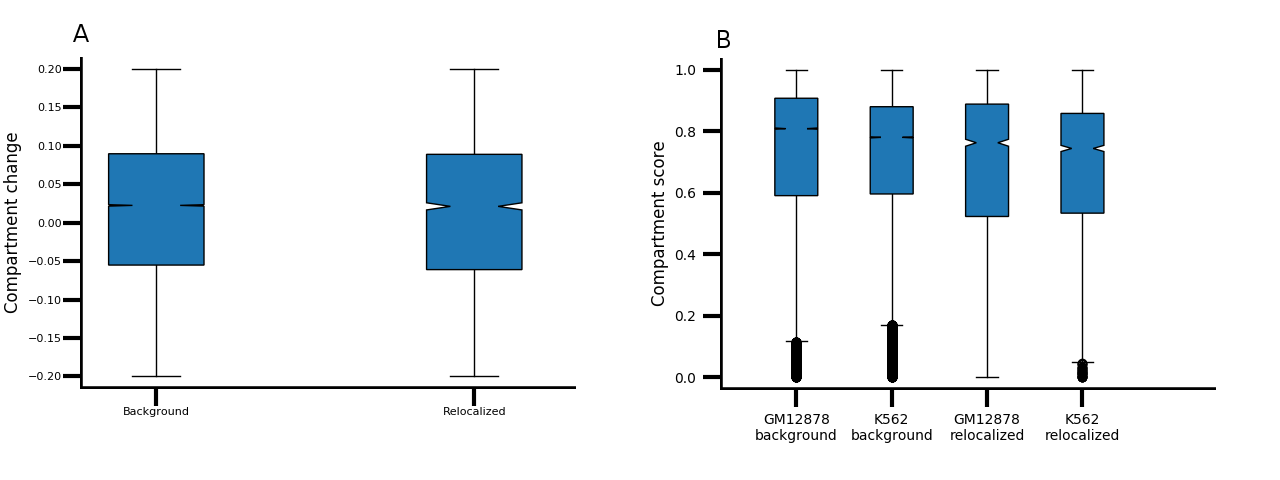

**Supplementary Figure 17.** Validation that compartment-independent relocalizations are not different from background in compartmentalization. **A)** Difference in compartment score in 10kb A compartment bins that did not significantly relocalize and compartment-independent relocalizations. **B)** Compartment scores for both cell types in 10kb A compartment bins that did not significantly relocalize and compartment-independent relocalizations.

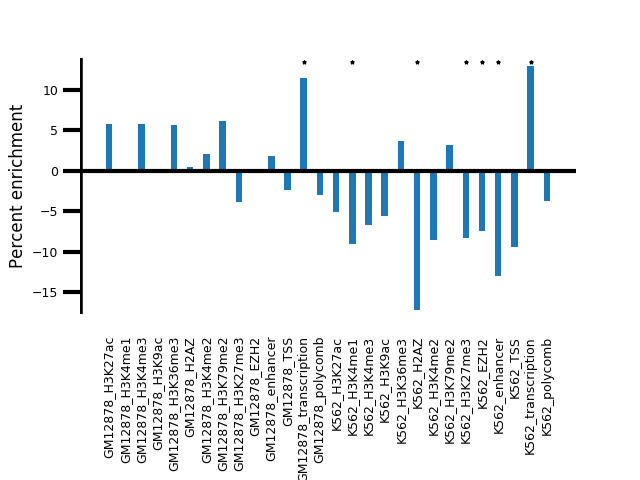

**Supplementary Figure 18.** Enrichment of mean coverage of selected chromatin marks in A compartment loci that relocalize without changing compartment score relative to those that do not relocalize, calculated from independent structural inference and alignment.

**Supplementary Table 1.** Compartment scores at chr21:47.4-47.5 Mb

| **Cell type** | **Compartment score** |
| --- | --- |
| IMR90 | 0.900 |
| HMEC | 0.756 |
| HUVEC | 0.767 |
| K562 | 0.853 |
